# Protein-coding potential of RNAs measured by potentially translated island scores

**DOI:** 10.1101/2021.04.14.439730

**Authors:** Yusuke Suenaga, Mamoru Kato, Momoko Nagai, Kazuma Nakatani, Hiroyuki Kogashi, Miho Kobatake, Takashi Makino

**Author notes:** These authors equally contributed to this work (Yusuke Suenaga, Mamoru Kato). **Correspondence to:** Yusuke Suenaga.

## Abstract

Recent studies have identified numerous RNAs that are functionally both coding and noncoding. However, the sequence characteristics that determine bifunctionality remain largely unknown. In this study, we developed and tested a potentially translated island (PTI) score, defined as the occupancy of the longest open reading frame (ORF) among all putative ORFs. We found that this score correlated with translation, including noncoding RNAs. In bacteria and archaea, coding and noncoding transcripts had narrow distributions of high and low PTI scores, respectively, whereas those of eukaryotes showed relatively broader distributions, with considerable overlap between coding and noncoding transcripts. The extent of overlap positively and negatively correlated with the mutation rates of genomes and effective population sizes of species, respectively. These overlaps were significantly increased in threatened species. In macroevolution, the appearance of the nucleus and multicellularity seem to have influenced the overlap of PTI score distributions, so that the probability of the existence of bifunctional RNAs is increased in eukaryotes. In mammalian testes, we observed an enrichment of noncoding RNAs with high PTI scores, which are candidates for bifunctional RNAs. These results suggest that the decrease in population size and the emergence of testes in eukaryotic multicellular organisms allow for the stable existence of bifunctional RNAs, consequently increasing the probability of the birth of novel coding and non-coding RNAs.

## Introduction

Recent advances in RNA sequencing technology have revealed that most of the eukaryotic genome is transcribed, primarily producing noncoding RNAs (Okazaki et al. 2002; Djebali et al. 2012; Ulitsky and Bartel 2013; Kopp et al. 2018). Noncoding RNAs that are more than 200 nucleotides in length are referred to as long noncoding RNAs (lncRNAs) and are not translated into proteins (Ulitsky and Bartel 2013; Kopp et al. 2018). LncRNAs have been reported to function in multiple biological phenomena, including the regulation of transcription, modulation of protein or RNA functions, and nuclear organization (Ulitsky and Bartel 2013; Kopp et al. 2018). However, paradoxically, a large fraction of lncRNAs are associated with ribosomes and are translated into peptides (Frith et al. 2006; Ingolia et al. 2011; Bazzini et al. 2014; Ingolia et al. 2014; Ruiz-Orera et al. 2014). Peptides translated from transcripts annotated as lncRNAs have been shown to have biological functions in multiple cases in eukaryotes (Li and Liu 2019; Huang et al. 2021), and some of these translations are specific to the cellular context (Dohka et al 2021). Conversely, known protein-coding genes, such as *TP53*, have second roles as functional RNAs (Candeias 2011; Kloc et al. 2011; Huang et al. 2021). The discovery of these RNAs with binary functions has blurred the distinction between coding and noncoding RNAs, so the characteristics of RNA sequences that explain the continuity between noncoding and coding transcripts remain unclear.

During evolution, new genes originate from pre-existing genes via gene duplication or from non-genic regions via the generation of new open reading frames (ORFs) (Ohno 1970; Chen et al. 2013; Zhang and Long 2014; McLysaght and Guerzoni 2015; McLysaght and Hurst 2016; Holland et al. 2017). The latter are *de novo* genes (Begun et al. 2006; Levine et al. 2006; Begun et al. 2007; Knowles and McLysagtht 2009; Li et al. 2009; Toll-Riera et al. 2009; Li et al. 2010), which have been shown to regulate phenotypes and diseases (McLysaght and Guerzoni 2015; Chen et al. 2013; Zhang and Long 2014), including brain function and carcinogenesis in humans (Li C-Y et al. 2010; Suenaga et al. 2014). lncRNAs serve as sources of newly evolving *de novo* genes (Ruiz-Orera et al. 2014), some of which encode proteins. In addition to ORFs exposed to natural selection, neutrally evolving ORFs are also translated from lncRNAs that stably express peptides (Ruiz-Orera et al. 2018), providing a foundation for the development of new functional peptides/proteins. High levels of lncRNA expression (Ruiz-Orera et al. 2018), hexamer frequencies of ORFs (Sun et al. 2013; Wang et al. 2013; Ruiz-Orera et al. 2014), and high peptide flexibility (Wilson et al. 2017) have been proposed as determinants of coding potential; however, the molecular mechanisms by which lncRNAs evolve into new coding transcripts remain unclear (Van Oss and Carvunis 2019).

In this study, we sought to identify a new indicator for determining RNA protein-coding potential. Within an RNA sequence, we defined sequence segments that start with AUG start codons and end with UAG, UGA, or UAA stop codons as potentially translated islands (PTIs). First, we defined this indicator using PTI lengths and subsequently examined the associations between the indicator and protein-coding potential. We also present analyses of more than 3.4 million transcripts in 100 organisms belonging to all three domains of life to investigate the relationship between the PTI score and protein-coding potential over evolutionary history.

## Results

### Coding transcripts show higher PTI scores in humans and mice

We previously identified a *de novo* gene, *NCYM*, and showed that its protein has a biochemical function (Suenaga et al 2014; Suenaga et al 2020). However, *NCYM* was previously registered as a non-coding RNA in the public database, and the established predictor for protein-coding potential (Wang et al 2013), the coding potential assessment tool (CPAT), showed a coding probability of NCYM of 0.022, labeling it as a noncoding RNA (Supplementary Figure 1). Therefore, we sought to identify a new indicator for coding potential, comparing *NCYM* with a small subset of coding and non-coding RNAs to determine whether NCYM has sequence features that would allow it to be registered as a coding transcript (data not shown). We found that predicted ORFs, other than major ORFs, seem to be short in coding RNAs. In addition, it has been reported that upstream ORFs inhibit the translation of major ORFs (Calvo et al 2009). Therefore, we hypothesized that the predicted ORFs may reduce the translation of major ORFs, thereby becoming short in the coding transcripts, including *NCYM,* during evolution. The term ORF refers to an RNA sequence that is translated into an actual product; however, the biological significance of non-translating, predicted ORFs has been largely ignored and remains to be characterized. Therefore, we defined a PTI as an RNA sequence from the start codon sequence to the end codon sequence and did not assume that it would result in a translated product. Thus, PTI can be defined even in genuine non-coding RNAs. The major ORFs are often the longest PTIs (hereafter, primary PTIs or pPTIs) in coding transcripts. Thus, to investigate the importance of pPTIs relative to other PTIs (hereafter, secondary PTIs, or sPTIs) for the evolution of coding genes, we defined a PTI score as the occupancy of the pPTI length to the total PTI length (Figure 1A–B) and assumed that the PTI score was high in coding transcripts. To examine this hypothesis, we first calculated the PTI scores for all human transcripts. We analyzed human transcripts from the National Center for Biotechnology Information (NCBI) nucleotide database for coding and noncoding (RefSeq accession numbers starting with NM and NR, respectively) transcripts. The data were downloaded using the Table browser (https://genome.ucsc.edu/cgi-bin/hgTables) after setting the track tab as “RefSeq Genes.” A total of 50,052 coding (NM) and 13,550 noncoding (NR) RNAs were registered in 2018 (Supplementary Table 1). To analyze putative lncRNAs with protein-coding potential, we excluded small RNAs (shorter than 200 bp) or RNAs with a short pPTI (shorter than 20 amino acids) from the NR transcripts, focusing on the remaining 12,827 transcripts.

**Figure 1.**
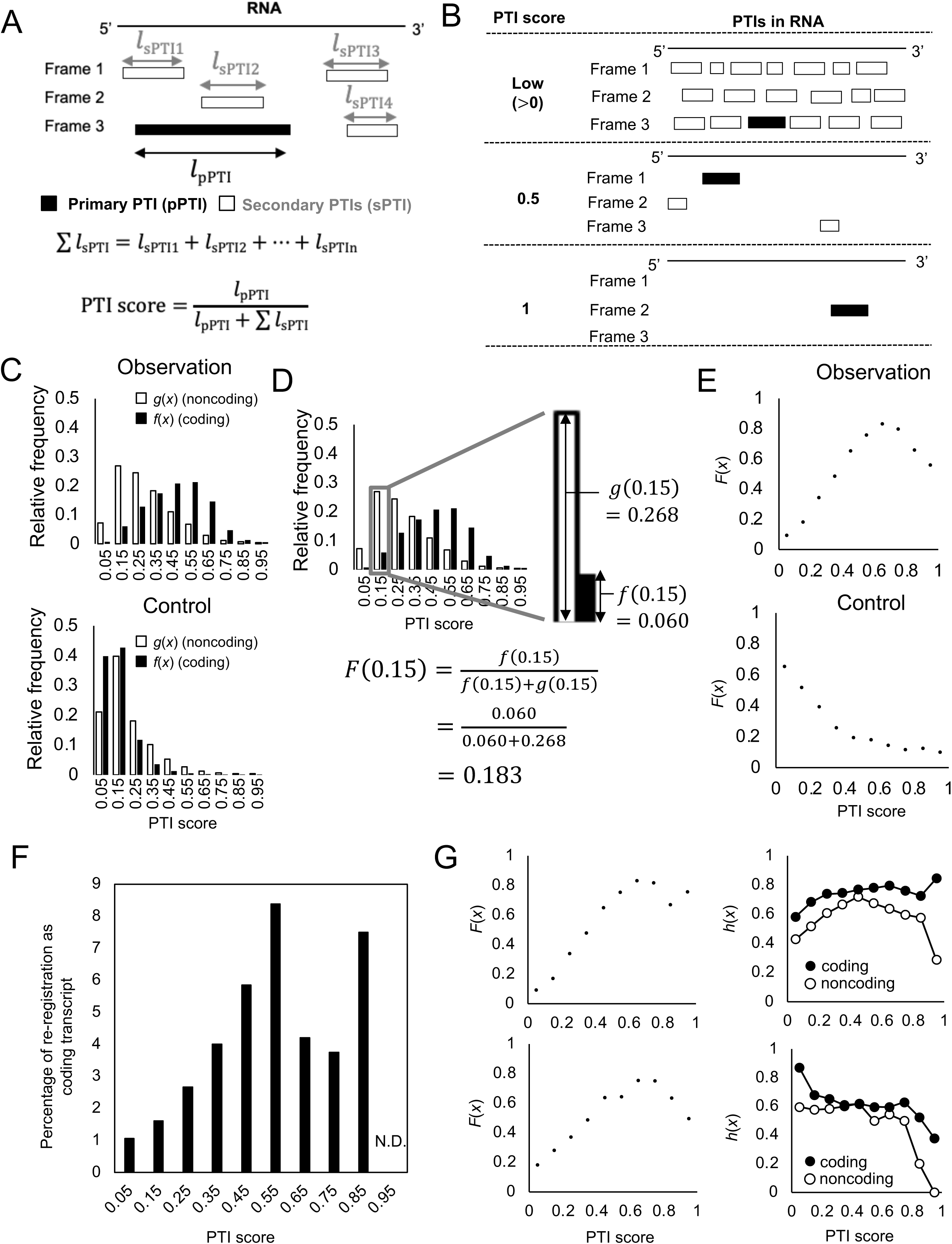
Potentially translated island (PTI) scores predict protein-coding potential of human transcripts. (A) Conceptual schematic of PTIs in an RNA in the three reading frames and definition of PTI score. Black and white rectangles indicate primary and secondary PTIs, respectively. The primary PTI is the longest PTI, while secondary PTIs are all others; *l* is PTI length. (B) Schematic of PTI distributions in RNAs with low (0-0.5), medium (0.5), and high (1) scores. (C) Relative frequencies of PTI scores of coding f(x) and noncoding g(x) transcripts (upper) and of random controls (bottom). (D) Explanation of *F*(x) for a PTI score of 0.15. (E) PTI score correlations with protein-coding potential, *F*(*x*), at PTI scores ≤ 0.65 (upper) and those in random controls (lower). (F) Relationship between PTI scores and percentages of NR transcripts re-registered as NM during the past 3 years. N.D., not detected. (G) Relationship between PTI scores and *F*(*x*) in human transcripts syntenic to chimpanzee (upper left) or mouse (bottom left). The relative frequency of transcripts with negative selection *h*(*x*) are plotted for each PTI score (upper and bottom right). The transcripts are syntenic to the genome of chimpanzee (upper right) or mouse (bottom right).The open circles indicate NR transcripts and the closed circles indicate NM transcripts.

We analyzed the relative frequencies of NM and NR transcripts, designated as *f*(*x*) and *g*(*x*), respectively (Figure 1C), where *x* indicates the PTI score. In human transcripts, *g*(*x*) showed a distribution that shifted to the left with an apex of 0.15; in contrast, the distribution of *f*(*x*) shifted to the right with an apex of 0.55 (Figure 1C, upper panel). As a control, we generated nucleic acid sequences in which A/T/G/C bases were randomly assigned with equal probabilities. In the controls, the relative frequencies of PTI scores were shifted to the left in both coding and noncoding transcripts (Figure 1C, bottom panel). The controls that randomly shuffled the original sequence without affecting the number of A/T/C/G bases in each transcript also had relative frequencies of PTI scores shifted to the left in both coding and noncoding transcripts (Supplementary Figure 2A). Similar results were obtained using a dataset from the Ensembl database (Supplementary Figure 2B). We also calculated the PTI scores of mouse transcripts from RefSeq and Ensembl and found that the distribution of *f*(*x*) was shifted to the right with an apex of 0.55 (Supplementary Figure 2C), similar to that of human transcripts. These results suggest that the sequences, not lengths, of the coding transcripts increased the PTI scores in mice and humans.

### PTI scores correlate with protein-coding potential in humans and mice

Next, we examined the relationship between PTI score and protein-coding potential. Based on the PTI score distributions of coding and noncoding transcripts, protein-coding potential *F*(*x*) was defined as the probability of being a coding transcript with a PTI score of *x*. A sample *F*(0.15) calculation for human transcripts is shown in Figure 1D. This result indicates that any given human RNA transcript with a calculated PTI score of 0.15 has a protein-coding potential *F*(*x*) of 0.183. *F*(*x*) was correlated with PTI scores ≤ 0.65 (Figure 1E and Supplementary Figure 3A). The protein-coding potentials of sequences in RefSeq data slightly decreased after peaking at 0.65 (Figure 1E), whereas those of sequences in the Ensembl data remained high (Supplementary Figure 3A). The *F*(*x*) of human transcripts was approximated by the following linear regression:

Based on Ensembl data,

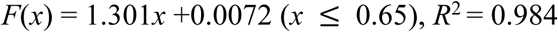

Based on RefSeq data,

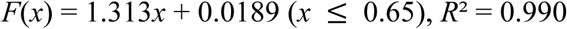

The intercepts were near zero, and the slopes were approximately 1.3. Using these formulas, we can calculate the protein-coding potential *F*(*x*) for any given human transcript with a PTI score of ≤ 0.65. For example, the *F*(*x*) of *NCYM* was calculated to be 0.746 and 0.765 based on the Ensembl and RefSeq databases, respectively (Supplementary Figure 1D). In contrast, *F*(*x*) for the controls was not correlated with the PTI scores (Figure 1E, bottom panel, and Supplementary Figure 3A). Similar results were obtained for the mouse transcripts (Supplementary Figures 3B). The *F*(*x*) of the mouse transcripts (PTI score ≤ 0.65) was approximated as follows:

Based on Ensembl data,

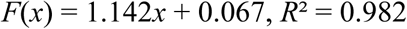

Based on RefSeq data,

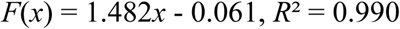

For both human and mouse transcripts, the PTI score correlated linearly with the protein-coding potential at PTI scores ≤ 0.65. Moreover, when the PTI score limit approached 0, the probability of the transcript being a coding RNA was 0 (Figure 1E and Supplementary Figure 3).

### Characterization of high-scoring human lncRNAs

Next, we investigated whether the PTI score is useful for identifying coding RNAs among NR transcripts. From the 7,144 transcripts registered as noncoding genes in 2015, we excluded small RNAs (< 200 nucleotides) and those with short primary PTIs (< 20 residues). Among the remaining 6,617 NR genes, 219 were reassigned as NM over the past 3 years (Supplementary Table 2), including the previously identified *de novo* gene *MYCNOS/NCYM* (Suenaga et al. 2014). The percentage of reclassification increased for NR transcripts with high PTI scores (Figure 1F). Thus, a high PTI score is a useful indicator of coding transcripts. NR transcripts with high protein-coding potential (0.6 ≤ PTI score < 0.8) were then extracted, and the domain structure of the pPTI amino-acid sequence was assessed using BLASTP. A total of 217 transcripts showed putative domain structures in pPTI, whereas 310 did not (Supplementary Table 3). Transcripts with domain structures are often derived from transcript variants, pseudogenes, or readthrough of coding genes; those without domain structures are often derived from antisense or long intergenic noncoding RNAs (lincRNAs) (Table 1).

**Table 1.** Numbers of original transcripts that produced NR transcripts with high coding

| Transcript | Domain |  | Total | P-value |
| --- | --- | --- | --- | --- |
|  | With | Without |  |  |
| Antisense | 4 | 61 | 65 | 7.79E-08 |
| LincRNA | 3 | 65 | 68 | 7.60E-09 |
| Pseudogene | 50 | 17 | 67 | 4.32E-07 |
| Readthrough | 7 | 0 | 7 | 6.00E-03 |
| Transcript variant of coding gene | 146 | 35 | 181 | 1.05E-19 |
| Divergent | 0 | 2 | 2 | N.S. |
| Intronic | 0 | 6 | 6 | N.S. |
| Small nuclear RNA | 0 | 3 | 3 | N.S. |
| miRNA host gene | 0 | 3 | 3 | N.S. |
| Other lncRNA | 7 | 118 | 125 | 1.12E-13 |
| Total | 217 | 310 | 527 |  |

*P*-values were calculated using the Yate's continuity correction. N.S., not significant.

We next examined the functions of genes generating NR transcripts with high coding potential (0.6 ≤ PTI score < 0.8). We divided the NR transcripts into those with and without putative domains to investigate novel coding gene candidates, either originating from pre-existing genes or created from non-genic regions. Analysis using the Database for Annotation, Visualization, and Integrated Discovery (DAVID) functional annotation tool (Huang et al. 2009a, 2009b) showed that NR transcripts without domain structures were derived from original genes related to transcriptional regulation, multicellular organismal processes, and developmental processes (Supplementary Table 4). Among the target genes of transcription factors, NMYC (MYCN), TGIF, and ZIC2 were ranked in the top three and are all necessary for forebrain development (Supplementary Table 4) (Brown et al. 1998; Gripp et al. 2000; van Bokhoven et al. 2005). We observed that NR transcripts with domain structures originating from genes that undergo alternative splicing are related to organelle function and are expressed in multiple cancers, including respiratory tract tumors, gastrointestinal tumors, retinoblastomas, and medulloblastomas (Supplementary Table 5). Similar analyses were conducted in mice (Supplementary Tables 6–8) and *C*. *elegans* (Supplementary Tables 9–11). In mice, the original genes related to protein dimerization activity (Supplementary Table 7) and nucleotide binding or organelle function (Supplementary Table 8) were enriched in high-PTI score lncRNAs with and without conserved domains, respectively. In *C*. *elegans*, the original genes related to embryo development (Supplementary Table 10) and chromosome V or single-organism cellular processes (Supplementary Table 11) were enriched. Therefore, the relationship between brain development and cancer in the function of high-PTI-score lncRNAs seems to be specific to humans.

### PTIs affect the protein-coding potential predicted by *K*a/*K*s

To examine the relationship between PTI scores and natural selection in the prediction of protein-coding potential, we calculated the ratio of nonsynonymous (*K*a) to synonymous (*K*s) values by comparing human transcripts with syntenic genomic regions of chimpanzees and mice (Figure 1G). Transcripts were selected based on the syntenically conserved regions: 44,593 (vs. chimp) and 14,016 (vs. mouse). We found a linear relationship between the *F*(*x*) and PTI scores in the conserved transcripts (Figure 1G, left panels). As predicted, coding transcripts exhibited *K*a/*K*s < 0.5 at a higher frequency than did noncoding transcripts, with large differences observed when for PTI scores > 0.9 or < 0.1, with the smallest difference for PTI scores of approximately 0.35 to 0.45 (Figure 1G, right panels). These results indicate that for transcripts with PTI scores near the highest or lowest values, the conservation of ORF/pPTI sequences (negative selection, *K*a/*K*s < 0.5) determines the coding potential. In contrast, for transcripts with PTI scores between 0.35 and 0.45, the conservation of ORF/pPTI sequences has almost no effect on the coding potential, and thus ORF/pPTI sequences have more potential to evolve neutrally. Therefore, noncoding transcripts showing both negative selection (*K*a/*K*s < 0.5) and the highest PTI scores may include new coding transcript candidates. We list 23 such transcripts in Supplementary Table 12, including four transcript variants of a previously identified lncRNA that encodes a tumor-suppressive small peptide, HOXB-AS3 (Huang et al 2017).

### Translation of small peptides shifts PTI score distributions

To investigate the effect of translation on the PTI score, we calculated the PTI scores of lincRNAs encoding small proteins and compared them with the PTI score distribution of all lincRNAs (Figure 2A). We found that lincRNAs translating small proteins shifted to higher PTI scores, and lincRNAs with PTI scores around 0.45 were increased compared to the distribution of all lincRNAs.

**Figure 2.**
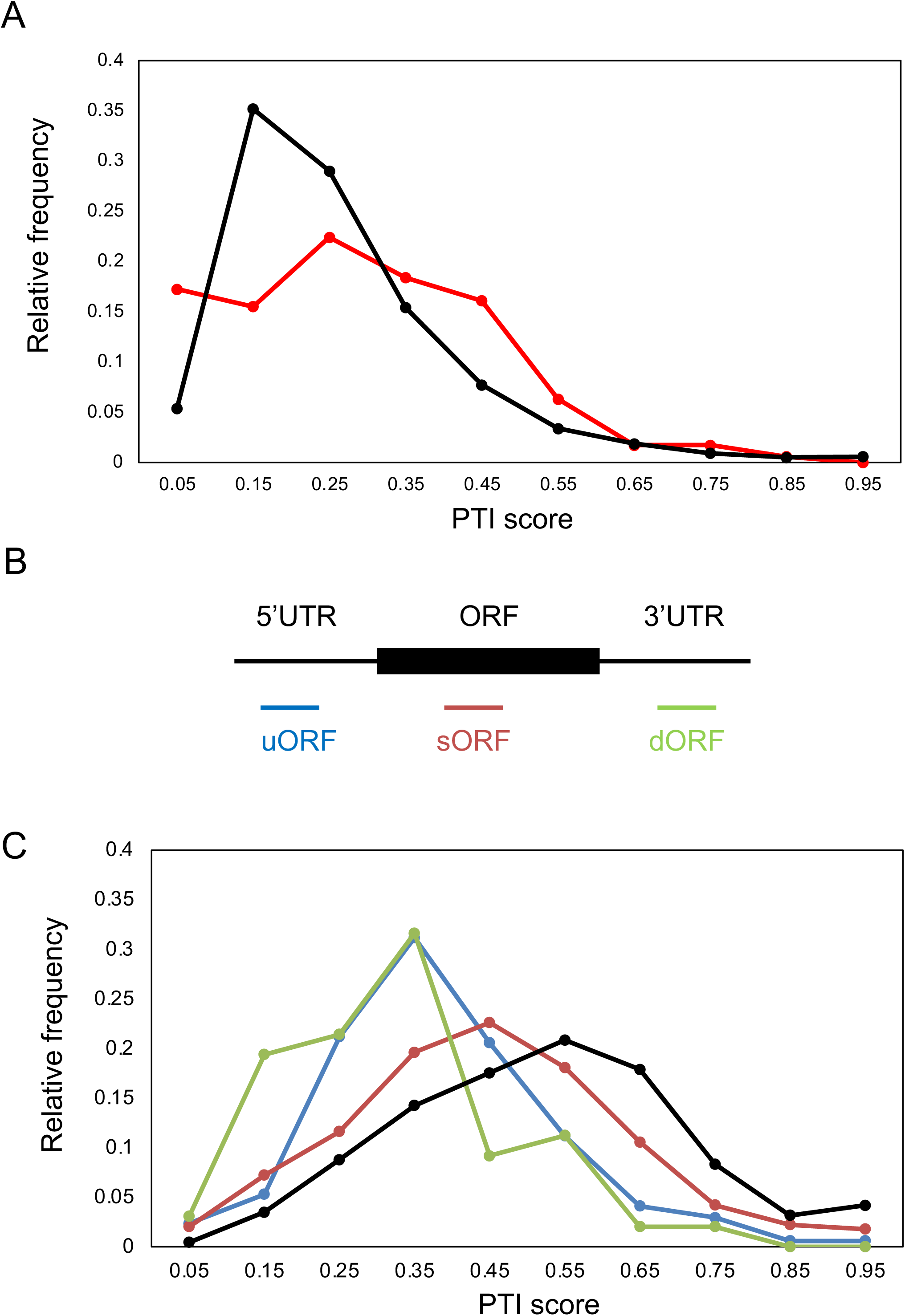
Translation effects on human PTI score distributions. (A) PTI score distribution of lincRNAs translating small proteins (red line, n = 174) registered in the SmProt database (http://bioinfo.ibp.ac.cn/SmProt/) shifts to higher scores relative to all lincRNAs registered in Ensembl (black line, n = 11,875). (B) Locations of uORF (blue), sORF(red), and dORF(green) relative to major ORF (black). (C) PTI score distributions of coding transcripts with translation of uORF (blue, n =170), sORF (red, n = 1,698), and dORF (green, n = 98) compared to all coding transcripts registered in Ensembl (black, n = 94,039).

ORF coverage, ORF size, and transcript length are indicators that have been used to predict the coding potential of transcripts (Wang et al. 2013; Zeng et al 2018). We calculated these three values for lincRNAs with translation products, and their distribution was compared with that of all lincRNAs. The comparison revealed no rightward shift in the peak, but there was a shift in the higher values of ORF coverage (Supplementary Figure 4A). On the other hand, there was no rightward shift in ORF size or transcript length (Supplementary Figure 4B and 4C). Therefore, the translation of non-coding RNAs was strongly correlated with the PTI score, but not with transcript length, ORF size, or ORF coverage.

Next, to examine whether PTI scores were associated with translation occupancy of ORFs in coding RNAs, we defined the ORFs in which translation products were identified in the sPTIs: uORFs, sPTIs with translation products detected in the 5’UTRs; sORFs, PTIs with translation products detected in other frames overlapping with ORFs of major proteins; and dORF, sPTIs with translation products detected in the 3’UTRs (Figure 2B). When the PTI scores of coding RNAs with uORF, sORF, and dORF were calculated and the PTI score distribution was compared with that of all coding RNAs, the PTI scores shifted to lower values, peaking at 0.35-0.45 (Figure 2C). Although there are differences in the effect of the location of the translated sPTI in a dataset from a different database, the PTI score distribution remained similar, that is, it shifted to lower values and increased the number of coding transcripts with PTI scores of 0.35-0.45 (Supplementary Figure 5). These results support the idea that the PTI score is related to the occupancy of major ORFs in the translation of RNAs. In addition, when considering the results in Figure 1G, translation of noncoding RNAs and sPTI in coding RNAs may increase the chances of pPTI/ORF sequences evolving neutrally by increasing transcripts with PTI scores of 0.35-0.45.

### Relationship between PTI score and relative frequencies of coding/noncoding transcripts in 100 organisms

To analyze the relationship between PTI scores and protein-coding potential in a broad lineage of organisms, we selected 100 organisms, consisting of five bacteria, ten archaea, and 85 eukaryotes (Supplementary Table 1), and calculated PTI scores for more than 3.4 million transcripts (Supplementary Table 1). Phylogenetic trees of the cellular organisms are presented on a logarithmic time scale, along with the number of species in each lineage used in the analyses in Figure 3. To examine the evolutionary conservation of the linear relationship between the PTI score and protein-coding potential in humans and mice, we selected a relatively large number of mammalian species (36). Species with fewer than three lncRNAs were not used to calculate *g*(*x*) and were not included in the histograms illustrating their relationship with the PTI score (Figures 4 and 5). For all organisms, the relative frequency of coding transcripts *f*(*x*) was shifted to the right (higher PTI score) compared to random or random shuffling controls (Figures 4 and 5; Supplement Figures 6 and 7).

**Figure 3.**
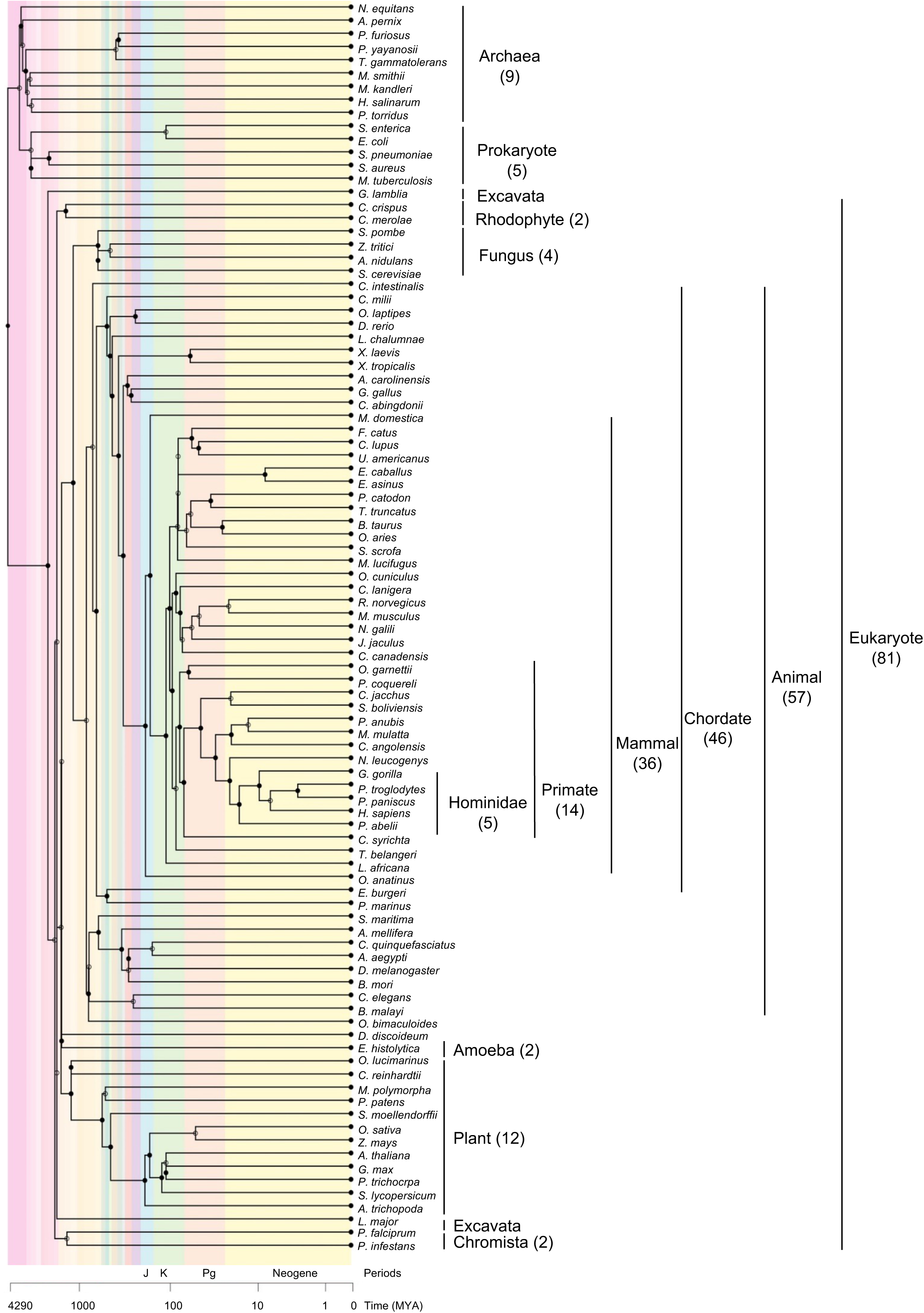
Phylogenetic tree. Numbers of species are indicated in each lineage. The lineages of five species, including one archaea (*Nitrososphaera viennensis* EN76), two fungi (*Puccinia graminis f. sp. Tritici* and *Pyricularia oryzae*), and two animals (*Strongylocentrotus purpuratus* and *Lingula anatine*) are unknown and excluded from the figure.

**Figure 4.**
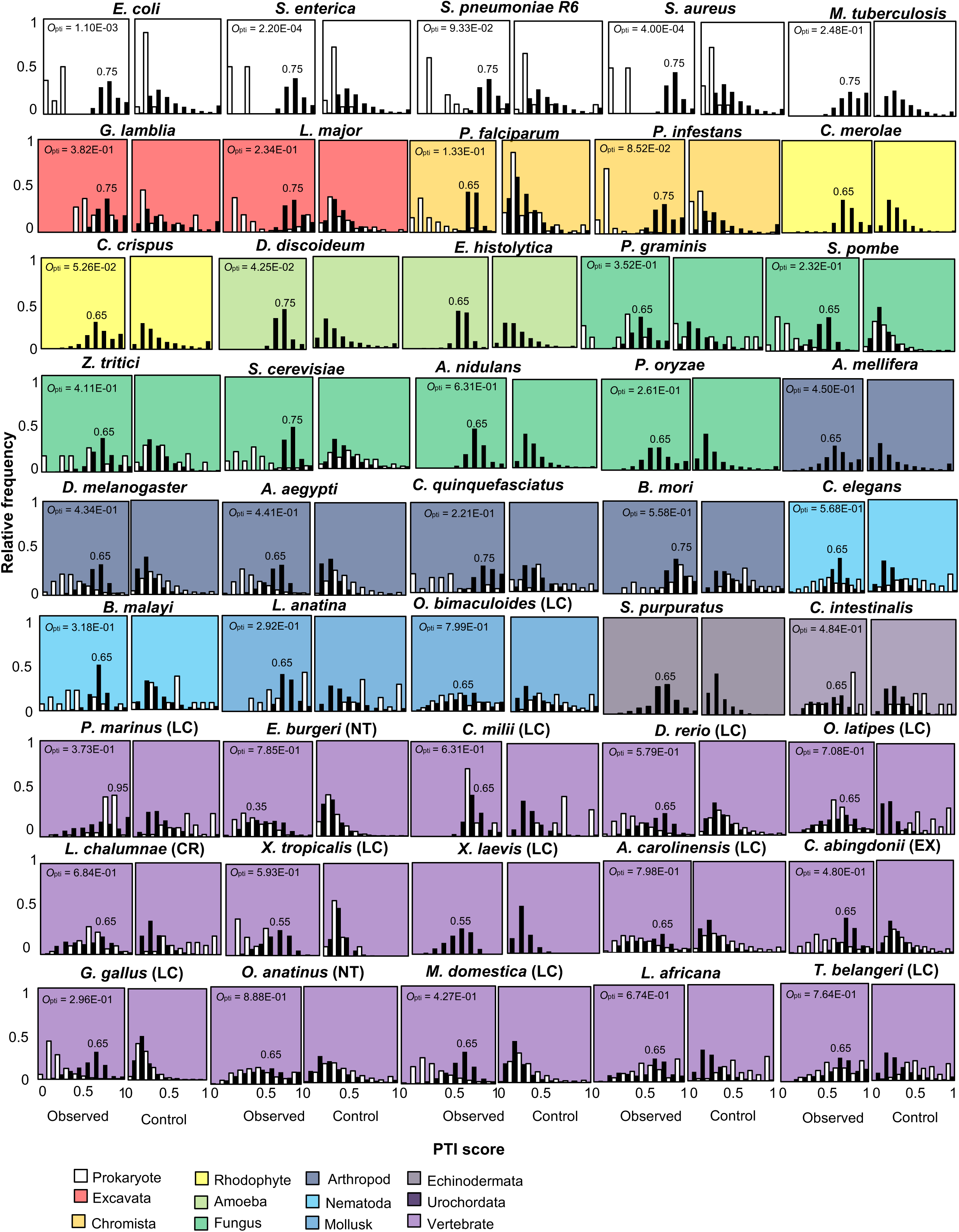
Relationships between PTI scores and relative frequencies of coding and noncoding transcripts from bacteria to mammals. Histograms of *f*(*x*) (white) or *g*(*x*) (black) in observed data (left) and in nucleic-acid–scrambled controls (right) for each species analyzed. PTI scores with the highest *f*(*x*) are presented in the histograms. *O*_pti_ was calculated using the PTI score distribution from observed data and indicated in the left panels.

**Figure 5.**
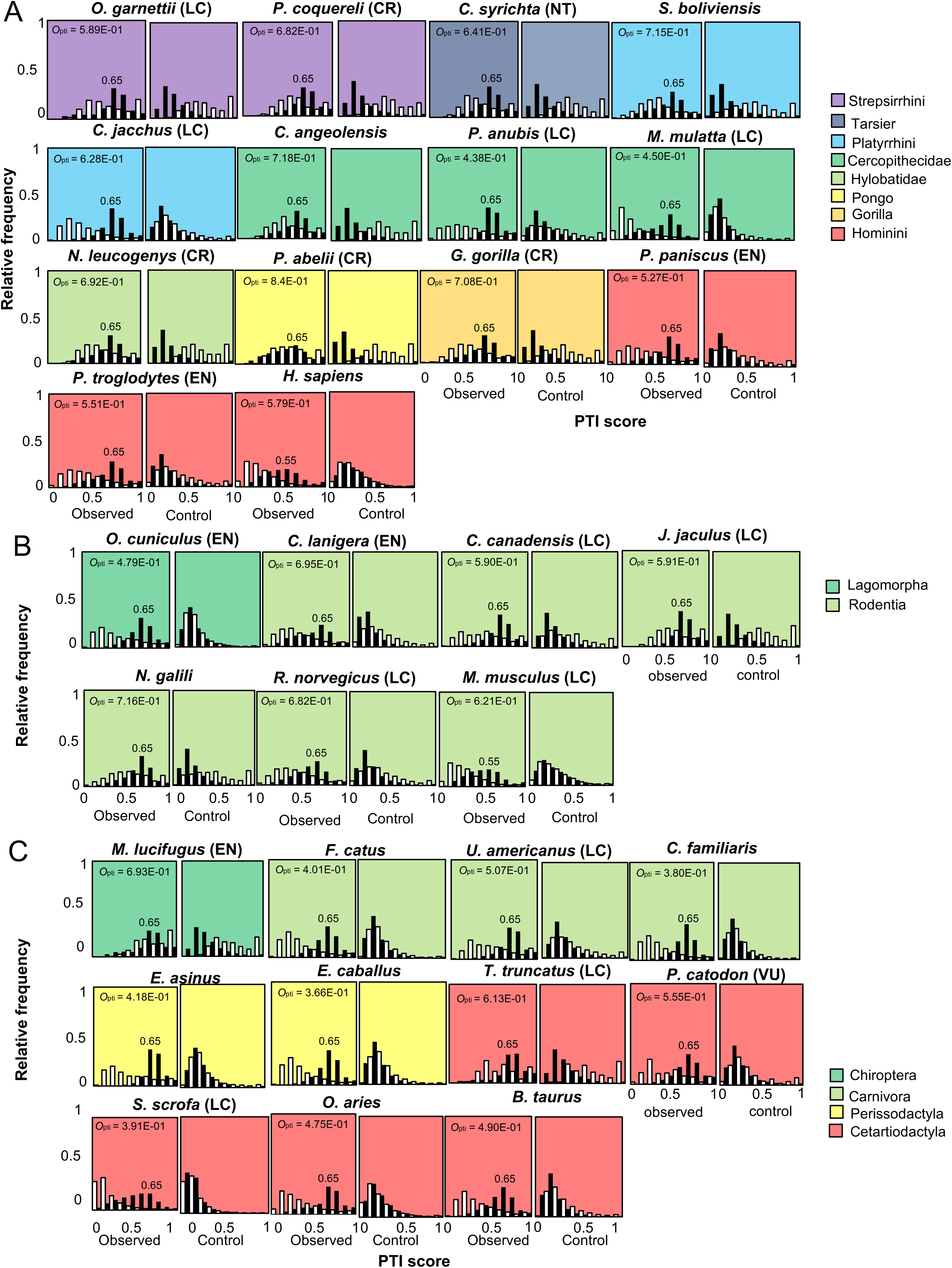
Relationships between PTI score and relative frequencies of coding f(x) and noncoding transcripts in Primates (A), Glires (B), and Laurasiatheria (C).

In bacteria and archaea, *f*(*x*) and *g*(*x*) exclusively exhibited high and low PTI scores, respectively, indicating a clear boundary between coding transcripts and lncRNAs in terms of PTI scores (Figure 4 and Supplementary Figure 6). In addition, the highest frequency of coding transcript *f*(*x*) presenting a PTI score was 0.75 in all examined bacteria (Figure 4) and ≥ 0.75 in archaea (Supplementary Figure 6). Among eukaryotes, unicellular organisms and non-vertebrates showed the highest frequencies of coding transcripts at 0.65 or 0.75 (Figure 4), while most vertebrates showed the highest values ≤ 0.65 (Figures 4 and 5). In addition, the *f*(*x*) distribution in vertebrates was broad and shifted to the left (lower PTI scores) relative to those of bacteria and archaea (Figure 4 and 5). In sharp contrast to *f*(*x*), the relative frequency of lncRNAs *g*(*x*) was shifted to the right (higher PTI scores) in eukaryotes, including *G. lamblia,* which belongs to the earliest diverging eukaryotic lineage and lacks mitochondria (Figure 4). Since the distribution of *f*(*x*) in the Excavata, including *G. lamblia,* showed a similar pattern to that of bacteria, the right shift of *g*(*x*) seems to be an earlier event than the left shift of *f*(*x*) in the evolution of eukaryotes. Collectively, the right and left shifts of *f*(*x*) and *g*(*x*) contribute to blurring the boundary between coding and noncoding transcripts in eukaryotes.

### PTI score distribution overlap is inversely correlated with effective population size

In general, eukaryotes (particularly multicellular organisms) have smaller effective population sizes than prokaryotes, with higher mutation rates due to the effect of genetic drift (Lynch et al 2016). We defined an indicator of coding/noncoding boundary ambiguity (overlapping score, *O*_pti_) and examined the relationship between *O*_pti_ and effective population size and mutation rate, using data from a previous study (Lynch et al 2016). The overlapping score based on ORF coverage, *O*_cov_, was also defined for comparison (Supplementary Figure 8). Of the 35 species used in this study, 11 had no more than five lncRNAs with pPTIs longer than 20 residues, and transcripts of the remaining 24 species (Supplementary Table 13 and Supplementary Figure 8) were used for the analysis. Similar to a previous report (Lynch et al 2016), the effective population size was inversely proportional to the mutation rate of genomic DNA, even in the remaining 24 species (exponent = −1.126, *R*_2_ = 0.6842, Figure 6A). *O*_pti_ positively and negatively correlated with mutation rates and effective population size, respectively, with relationships that could be approximated as logarithmic (*R*_2_ = 0.7578) or exponential functions (*R*² = 0.4667). In contrast, ORF coverage (*O*_cov_) showed a weaker relationship with mutation rates and effective population size (Supplementary Figure 9). Substituting the maximum value of *O*_pti_, 1 into this exponential approximation (Figure 6A, right upper panel) yields the minimum effective population size, which is 1001.28. This is consistent with the observation that the minimum effective population size in conservation biology is approximately 1000 (Frankham et al 2014). This result led us to consider the possibility that *O*_pti_ might be elevated in endangered organisms. We calculated *O*_pti_ for 35 vertebrate species on the IUCN Red List (left panel, Figure 6B; Supplementary table 1), and found that species at risk of extinction had significantly higher *O*_pti_ than species with little risk of extinction (Least Concern, LC). In addition, among LCs, *O*_pti_ was higher for species with decreasing numbers compared to those with stable populations (right panel, Figure 6B; Supplementary table 1).

**Figure 6.**
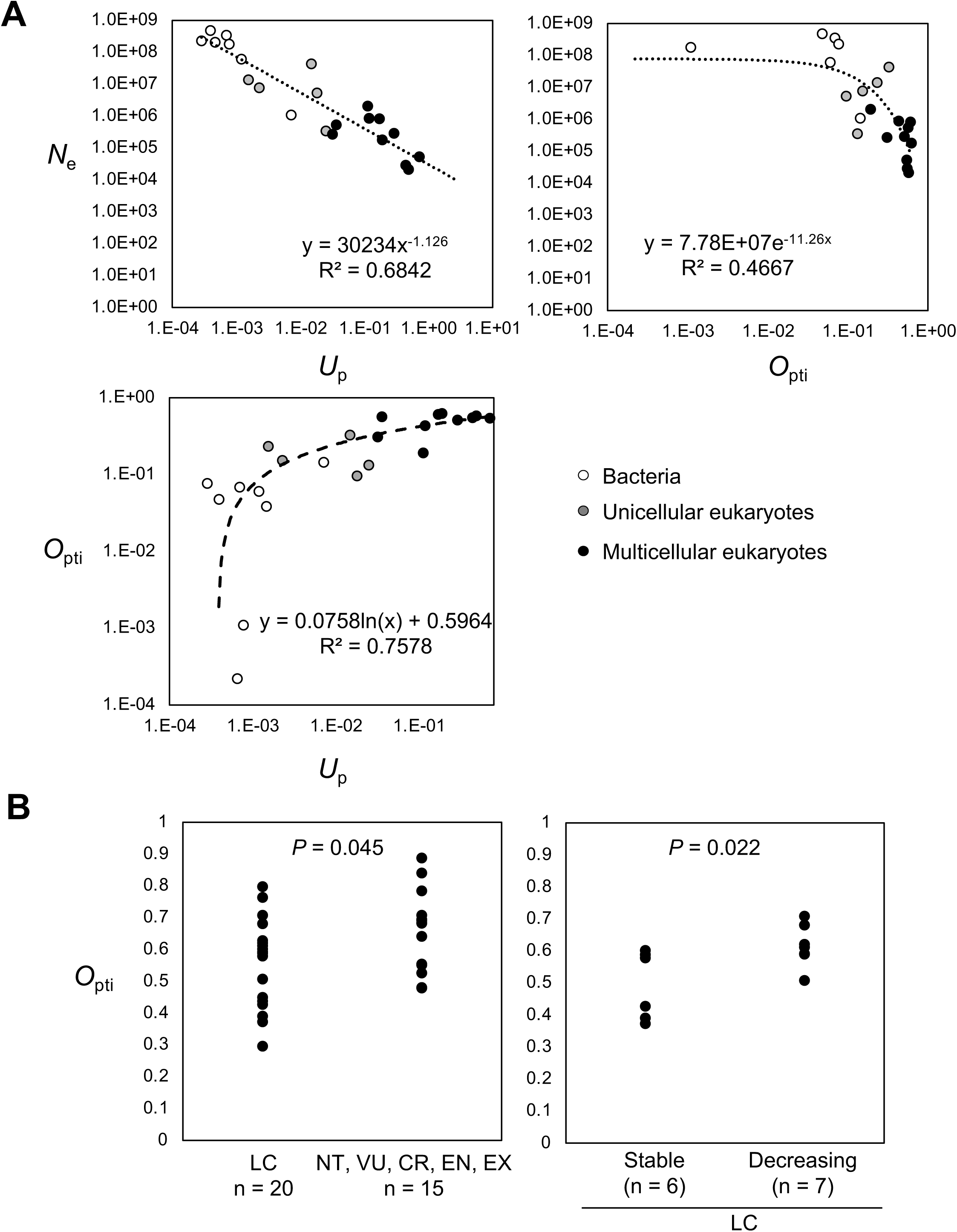
Overlap of PTI score distributions is negatively correlated with effective population sizes. (A) Inversely proportional relationship between genome-wide mutation rates in protein-coding DNA per generation (*U*_p_) and effective population sizes (*N*_e_) in 24 species (left upper). Values are from Lynch et al., 2016. *O*_pti_ positively and negatively correlates with *U*_p_ (left bottom) and *N*_e_ (right upper); these relationships are approximated by logaritic and exponential functions, respectively. White, gray, and black dots indicate bacteria, unicellular eukaryotes, and multicellular eukaryotes, respectively. (B) *O*_pti_ is increased in vertebrates at risk of extinction (left) and with decreasing population trends (right). LC, least concern (n = 20); NT, near threatened (n = 3); VU, vulnerable (n = 1); EN, endangered (n = 5); CR, critically endangered (n = 5); EX, extinct (n = 1). *P*-values were calculated by the Mann–Whitney U test.

### Relationship between PTI score and protein-coding potential

The overlapping of relative frequencies in *f*(*x*) and *g*(*x*) led us to examine the relationship between the PTI score and protein-coding potential *F*(*x*) in eukaryotes. To avoid being misled by small sample numbers, we selected 32 species with more than 1000 lncRNAs that contained pPTIs to calculate *F*(*x*) (Figure 7 and Supplementary Figure 10). In humans and mice, the relationship between the PTI score and *F*(*x*) was approximated with a linear function passing through the origin of the PTI score ≤ 0.65. Therefore, we used linear approximation of the *F*(*x*) of 32 species and found that 27 of the 32 species were well approximated by linear functions (indicated as linear group, L, in Figure 7 and Supplementary Figure 10). In *U. americanus*, *C. canadensis*, and *G. gorilla*, fewer than five lncRNAs exhibited PTI scores of 0.05; thus, we eliminated the *F*(0.05) in these species for the approximation by linear function (indicated with asterisks in Figure 7). The *F*(*x*) of the remaining five species that showed *O*_pti_ > 0.7 did not fit in linear approximations (indicated as constant group C in Figure 7) and were characterized by low slope values. They belonged to plants (*Z. mays*), reptiles (*A. carolinensis*), and mammals (*O. anatinu*, *S. boliviensis*, and *G. gorilla*) (Figure 7). In these species, PTI scores showed a weaker association with protein-coding potential. We noticed that these species may have small effective population sizes, possibly because of the risk of extinction (*O. anatinu* and *G. gorilla*) or artificial selection as pets (*A. carolinensis* and *S. boliviensis*) or as crops (*Z. mays*).

**Figure 7.**
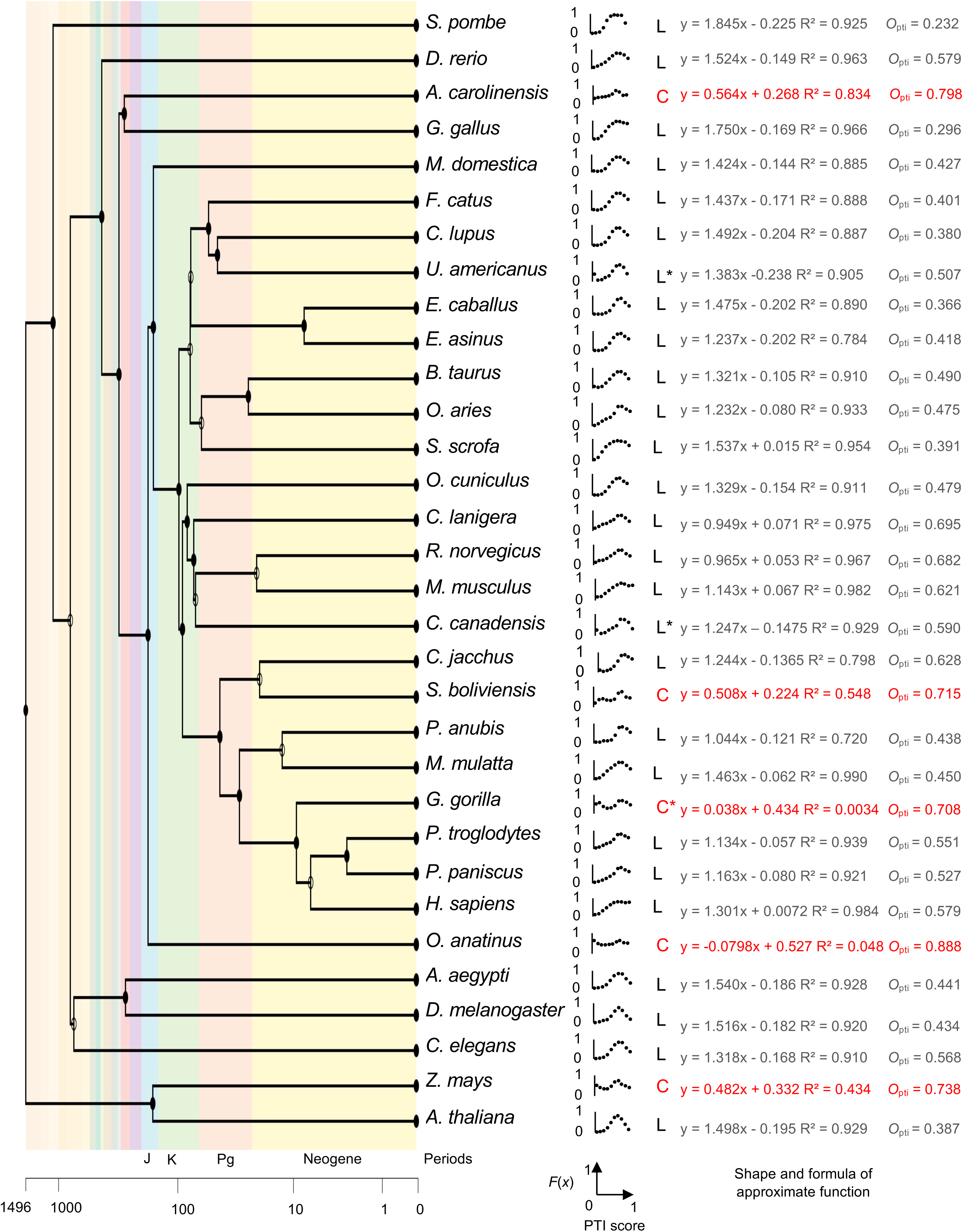
Relationship between PTI score and protein-coding potential *F*(*x*) for 32 eukaryotes. Phylogenetic tree including the 32 species (left), dot plots, and shape and formulas of approximate functions. L and C indicate linear (in black) and constant (in red) functions. Fewer than five lncRNAs had a PTI score of 0.05 in *U. americanus*, *C. canadensis*, and *G. gorilla*; therefore, we eliminated the *F*(0.05) for these species for linear function approximations (asterisks). *O*_pti_ was calculated using PTI score distributions of observed data.

### Characteristics of RNA virus genomes in human and bacterial cells

In sharp contrast to the coding transcripts of bacteria and archaea, the PTI scores of coding transcripts of eukaryotes overlapped with those of noncoding RNAs due to their broad distribution of low PTI scores. To investigate the molecular mechanism underlying the distinct distribution of coding transcripts between bacteria and eukaryotes, we analyzed the genome sequences of RNA viruses that infect human or bacterial cells. Positive-sense single-stranded RNAs, or (+) ssRNAs, are parts of the viral genome that generate mRNAs and are translated into viral proteins via the host translation system. Therefore, efficient translation in host cells contributes to the replication of (+) ssRNA viruses. We speculated that PTIs other than bona fide ORFs affect the coding potential of the viral genome in host cells. Multiple bona fide ORFs are present in viral genomes. Thus, we extended the concept of PTIs to multiple ORFs in viral RNA genomes (Figure 8A) and set the viral ORF (vORF) score.

**Figure 8.**
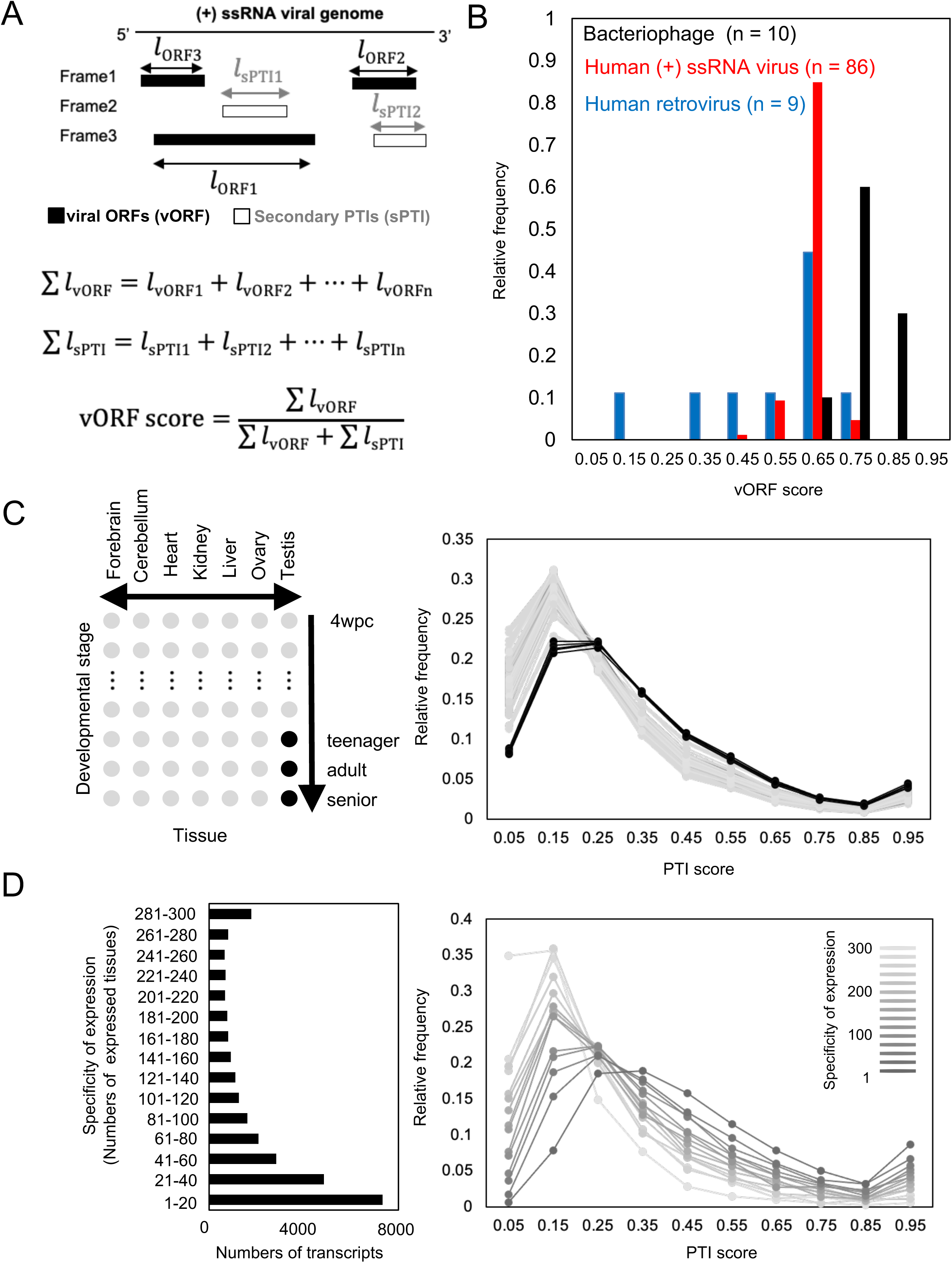
Molecular mechanisms that affect PTI score distributions. (A) Schematic explanation of sPTI length and bona fide viral ORFs in a (+) ssRNA virus genome and the definition of viral ORF (vORF) score. Black and white rectangles indicate viral ORFs and secondary PTIs, respectively. *l* is the length of the ORFs and PTIs. (B) Histograms of relative frequencies of human (+) ssRNA viruses (red) and bacteriophages (black). (C) PTI score distributions of lncRNAs in human tissues. Distributions in mature testes and other tissues are indicated as black and gray lines, respectively. (D) The relationship between tissue-specificity and PTI score distributions in humans. Line intensity represent specificity of gene expression.

Among the positive sense ssRNA viruses registered in the NCBI database, 198 were human viruses and 13 were bacteriophages. We eliminated the viruses that produced viral proteins by exceptional translation mechanisms such as ribosome frameshifting, alternative initiation sites, ribosome slippage, and RNA editing, focusing on the remaining 95 human viruses, including nine retroviruses (Supplementary Table 14) and 10 bacteriophages (Supplementary Table 15). The relative frequencies of the human viruses and bacteriophages showed distinct peaks at PTI scores of 0.65 and 0.75, respectively (Figure 8B). These values correspond to the PTI scores of the highest protein-coding potential in humans (Figure 1E and Supplementary Figure 3A) and the highest frequency of coding transcripts in bacteria (Figure 4). In addition, the relative frequency of human viruses showed a broader distribution of low PTI scores compared to bacteriophages, particularly in human retroviruses (Figure 8B). Therefore, RNA virus genomes appear to have sequence characteristics that maximize their protein-coding potential in host cells.

### The relationship between PTI scores and tissue-specific expressions

The right shift of the PTI score distribution for noncoding RNAs is pronounced in eukaryotes, especially in multicellular organisms (Figures 4 and 5). To examine the possibility that different tissues in multicellular organisms show different PTI distributions for noncoding RNAs, we analyzed transcriptome data to calculate the PTI scores of human noncoding transcripts expressed in multiple tissues (Figure 8C). The PTI score distributions were similar for almost all tissues, but, as an exception, they shifted to higher values for mature testes (Figure 8C). Similar results were also obtained for opossums, rats, mice, and macaques, although their shifts were weaker than those of humans (Supplementary figure 11). Furthermore, the noncoding transcripts that were expressed in a tissue-specific manner had higher PTI scores than ubiquitously expressed noncoding transcripts in humans (Figure 8D) and the other four species (Supplementary figure 12). The relationship between the specificity of expression and the PTI score was also found for human coding transcripts (Supplementary figure 13). These results suggest that the tissue-/cell type-specific expression of transcripts evolved in multicellular eukaryotes contributes to increased PTI scores for noncoding transcripts. Since the majority of tissue-specific transcripts were expressed in matured testes (7,573 of 8,523 transcripts (89%) in the highest specificity group for humans), the evolution of the testis also seems to contribute to the existence of high PTI score-noncoding RNAs, thus contributing to the birth of new coding genes.

## Discussion

Here, we showed that PTI scores are associated with protein-coding potential in cellular organisms. In bacteria and archaea, the PTI-score distributions for noncoding and coding transcripts were distinct (low and high scores), whereas they were merged in eukaryotes.

Right shifts in the distribution of noncoding RNA occurred in *G. lamblia*, one of the earliest diverging eukaryotes, which are binucleate and lack mitochondria, peroxisomes, and a typical Golgi apparatus (Ankarklev et al. 2010; Bartelt et al. 2015; Buret et al. 2020) and were commonly observed in all eukaryotes examined. Moreover, functional noncoding RNAs, by definition, should not be translated by ribosomes in cells. However, in bacteria and archaea, newly transcribed RNAs are immediately bound by ribosomes (Miller et al. 1970; French et al. 2007) and do not have the chance to escape translation. Thus, as expected, transcripts with noncoding functions in bacteria and archaea showed low PTI scores (top panel, Figure 9A). Alternatively, in eukaryotes, the existence of the nucleus prevents the immediate binding of lncRNAs by ribosomes, so cytoplasmic translocation from the nucleus is required for translation. Therefore, eukaryotic lncRNAs may function in the nucleus even with high PTI scores, and the subsequent evolution of cytosolic translocation of these noncoding RNAs may contribute to the origination of new coding genes (middle panel, Figure 9A). Thus, the pervasive transcription of the genome seems to help eukaryotes to create new functional noncoding/coding RNAs, while being disadvantageous for bacteria and archaea by increasing the risk of transcription of high-PTI-score transcripts, leading to immediate translation of wasteful and/or toxic proteins (top and middle panels, Figure 9A, Monsellier et al. 2007). In addition, multicellular organisms have a variety of intracellular environments because of the large number of cell types, which may increase the possibility of the existence of an intracellular environment in which newly created proteins are not toxic (bottom panel, Figure 9A). Since the possibility that a new protein will not be toxic in multiple intracellular environments is lower than the possibility that it will not be toxic in a particular intracellular environment, noncoding RNAs that are ubiquitously expressed need to have lower PTI scores than those with specific expression (bottom panel, Figure 9A).

**Figure 9.**
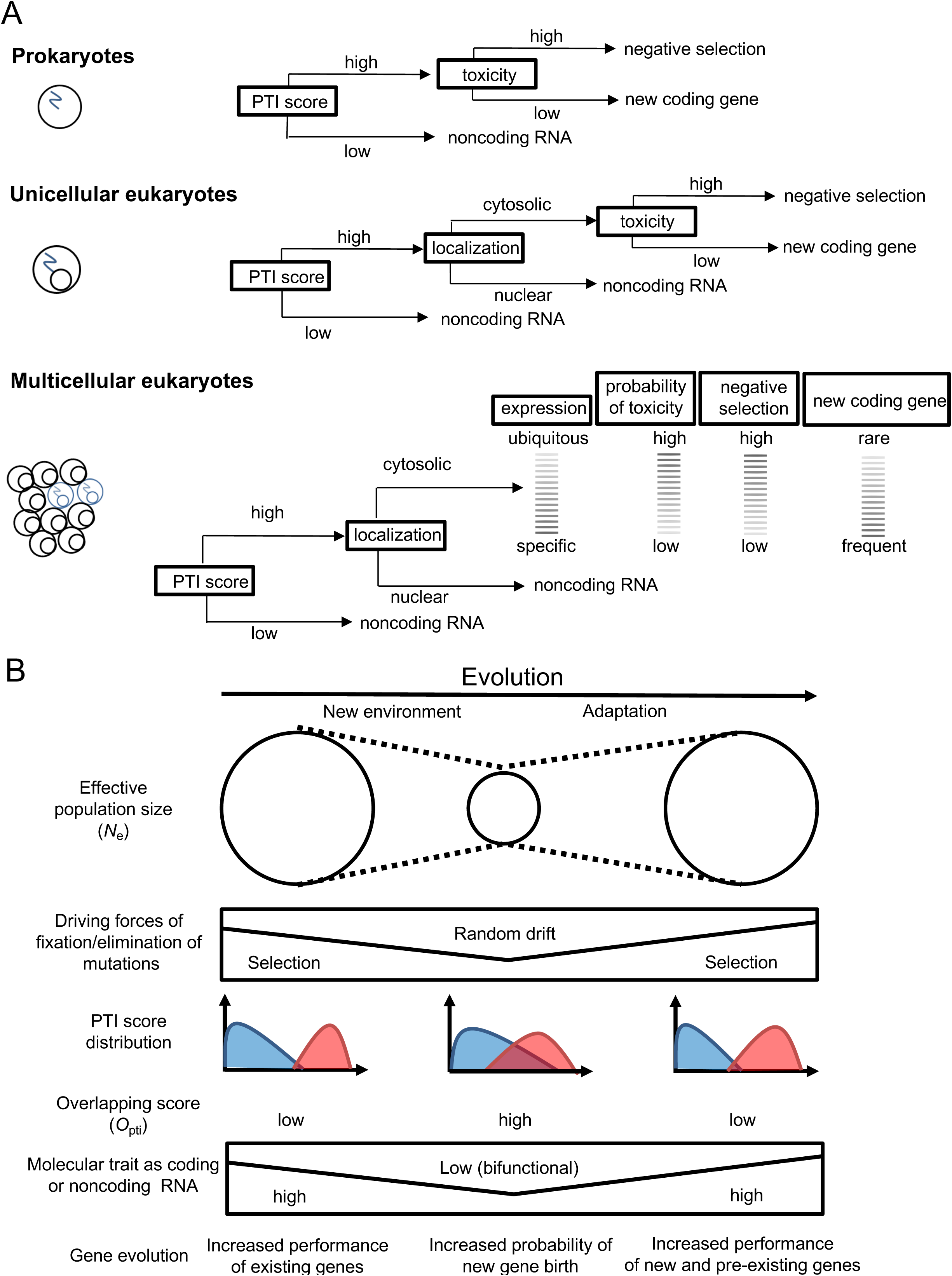
Hypothesis: new gene birth is a countermeasure to decline in effective population size. (A) Schematic explaining how nuclear evolution and multicellularity contribute to the generation of noncoding RNAs with high PTI scores in eukaryotes. (B) Schematic illustrating new gene birth in response to decline in effective population size caused by environmental changes.

Kaessmann proposed an “out of the testes hypothesis,” arguing that the testis facilitates the birth and evolution of new genes in animals. The germ cells (spermatocytes and spermatids) in the testes have an active chromatin state, and global transcription occurs, increasing the possibility of generating new coding genes (Kaessmann 2010). Consistent with this hypothesis, our results showed that the PTI score distribution of noncoding RNAs shifted to higher values only in mature testes with spermatocytes and spermatids, but not in immature testes or in other tissues. In addition, most transcripts with tissue-specific expression were found in mature testes, and these noncoding RNAs had high PTI scores. These results suggest that new coding genes are generated from noncoding RNAs with high PTI scores that are specifically expressed in germ cells of the mature testis.

Multiple human noncoding RNAs with high PTI scores have been reclassified as coding genes over the past 3 years, including the human *de novo* gene *NCYM* (Suenaga et al. 2014; Suenaga et al. 2020). Because *de novo* gene products have no known domain structures, high PTI score-noncoding transcripts without putative domains may be good candidates as novel *de novo* genes in eukaryotes. *NCYM* is an antisense gene of *MYCN*, whose protein product stabilizes the MYCN protein (Suenaga et al. 2014). MYCN directly stimulates *NCYM* and *OCT4* transcription, whereas OCT4 induces MYCN (Kaneko et al. 2015). This functional interplay forms a positive feedback loop, allowing these genes to induce each other’s expression in human neuroblastomas (Suenaga et al. 2014; Islam et al. 2015; Kaneko et al. 2015; Shoji et al. 2015). Functional annotation of noncoding genes without putative domains was related to transcriptional regulation, and the target genes of transcription factors, including MYCN, TGIF, and ZIC2, were enriched. As *de novo* emergence of *NCYM* occurred in Homininae, NCYM-mediated MYCN activation may modulate human *de novo* gene births during evolution, regulating the transcription of *MYCN* target genes. Notably, both NCYM and MYCN are expressed in germ cells of the testes (Suenaga et al 2014; Kanatsu-Shinohara et al 2016), and MYCN has been shown to regulate the self-renewal of spermatogonial stem cells (Kanatsu-Shinohara et al 2016). Furthermore, a recent study showed that binding sites for transcription factors, including MYCN, are mutational hotspots in human spermatogonia (Kaiser et al. 2021). Both *TGIF* and *ZIC2* are mutated in holoprosencephaly, a disorder caused by a failure in embryonic forebrain development (Brown et al. 1998; Gripp et al. 2000), whereas *MYCN* mutations cause Feingold and megalocephaly syndromes, which are associated with reduced and increased brain size, respectively (van Bokhoven et al. 2005; Kato et al. 2019). Thus, the present study also provides a list of candidate human *de novo* genes possibly involved in brain development and brain-related diseases.

Relative frequencies of positive-sense ssRNA viruses exhibit sharp peaks at vORF scores of 0.75 in bacteriophages and 0.65 in human viruses, indicating the adaptation of RNA viruses to host cells by maximizing the protein-coding potential of their genomes. Immediately after viral infection, the viral (+) ssRNA genomes, save for those of retroviruses, are used as templates for translation in the host cytosol. Thus, the distinct translation systems between humans and bacteria likely affect the left shift in the viral genome peak, as well as the left shift in coding RNA distribution in eukaryotes. In retroviruses, reverse transcriptase produces double-stranded DNA using the viral genome as a template, which is then inserted into the host genome. The viral genome is subsequently transcribed in the nucleus, and its mRNA is transported to the cytoplasm where protein products are translated in a manner similar to that of host proteins. Therefore, the relatively lower vORF score distribution in human retrovirus genomes is likely a function by the nuclear localization of the provirus, which may promote the diversification of PTI scores of RNA genomes via adaptation to host cellular mechanisms other than translation, such as cytosolic translocation. While the overlap of PTI score distributions of coding/noncoding transcripts seems to be beneficial by facilitating new gene birth, excessive overlap in PTI score distributions was found in species at the risk of extinction. Because the extent of the overlap (*O*_pti_) positively and negatively correlates with higher mutation rates and effective population sizes, respectively, the small effective population sizes in multicellular eukaryotes seem to increase the overlap by accumulation of the slightly deleterious or beneficial mutations driven by random drift, as predicted by neutral theory (Kimura 1968; Kimura 1983) and nearly neutral theory (Ohta 1992). Translation of noncoding RNA or sPTIs in coding RNAs caused right and left shifts in PTI score distributions, respectively. The mutations that cause these translations may be beneficial for increasing the possibility of evolution of new functional RNAs or regulatory mechanisms and be deleterious for inhibiting existing coding/noncoding functions. Translation of small proteins from noncoding RNAs seems to inhibit the noncoding functions of RNA because of ribosome binding and subsequent translation. In contrast, translation of sPTIs in coding RNAs seems to inhibit translation of major ORFs because of the competition for translation (Calvo et al 2009) without the evolution of specific regulatory mechanisms, such as the recently discovered mechanism in dORF (Wu et al. 2021).

According to the drift-barrier hypothesis (Lynch et al 2010; Lynch et al 2016), the performance of any molecular trait is expected to become more refined in larger population sizes, because the effects of selection relative to random drift are stronger. Consistent with this hypothesis, we found that the molecular traits of coding or noncoding RNAs were prominent in bacteria/archaea and weak in multicellular eukaryotes, allowing the existence of bifunctional RNAs. The excessive overlap of PTI score distributions (*O*_pti_ > 0.7) diminished the correlation between the PTI score and protein-coding potential. This indicates that both coding and noncoding transcripts lost their molecular traits as coding and noncoding RNAs in terms of PTI score, which became lethal or highly deleterious for the species.

Species with decreasing population sizes showed significantly higher *O*_pti_ compared with species with a stable population size, even in the LC group. Combined with the results discussed above, we propose a novel model of new gene origination in which new gene birth occurs in response to decreased effective population sizes (Figure 9B). At stable population sizes, natural selection maintains molecular traits of existing genes, and thus existing coding and noncoding functions of RNA stably exist with high and low PTI scores with low overlap of PTI score distribution of coding/noncoding transcripts. When new environments reduce the effective population size of species, the driving force of fixation/elimination of mutations changes from natural selection to random drift. This increases the probability of fixation of slightly deleterious/beneficial mutations, resulting in an increase in the overlap of PTI score distributions between coding and noncoding transcripts. The overlap allows the existence of bifunctional RNAs as candidates for new functional coding or noncoding transcripts. When the effective population size approaches 1,000 because of rapid decline, the accumulation of deleterious mutations decreases the long-term evolutionary potential of populations, leading to extinction. On the other hand, when the speed is slow enough for beneficial mutations to be fixed in the populations, the newly evolved coding/noncoding transcripts contribute to an increase in the effective population size, resulting in adaptation of the species to new environments. The increase in the effective population size leads to an increase in the effect of natural selection on the new functions of coding/noncoding genes as well as pre-existing genes. Thus, if nuclear evolution and multicellularity contribute to the generation of lncRNAs with high PTI scores and subsequent generation of novel coding genes (Figure 9A), the ability to generate new genes in response to population decline (Figure 9B) may be greatest in eukaryotic multicellular organisms.

In conclusion, the PTI score is an important indicator for integrating the concept of gene birth into classical evolutionary theory, thereby contributing to the elucidation of the molecular basis for the evolution of complex species, including humans. In the future, it will be necessary to calculate PTI scores based on the transcriptomes of additional species to test our hypothesis that positioning new gene birth as a countermeasure to the decline in effective population size.

## Materials and Methods

### Potentially translated islands (PTIs)

#### Definition

PTIs are defined as sequence segments beginning at AUG and ending with any of the UAA, UAG, or UGA stop codons in the 5ʹ to 3ʹ direction within an RNA sequence in all three possible reading frames (Figure 1A).

#### Example

The PTIs in the human *de novo* gene *NCYM* (Suenaga et al. 2014) were identified using the cDNA sequence (Supplementary Figure 1A) and are shown in bold characters (Supplementary Figure 1B). Further information is included in the Supplementary Notes.

### The length of a PTI and primary/secondary PTIs

#### Definition

The PTI length is defined as the length of the amino acid sequence, excluding the stop codon, and is represented by *l* (Figure 1A). In an RNA sequence, the longest PTI is designated as the primary PTI (pPTI), whereas the others are termed secondary PTIs (sPTIs). The lengths of pPTI and sPTI are described as *l*_pPTI_ and *l*_pPTI_, respectively (Figure 1A).

#### Example

The shortest possible PTI is “AUGUAA,” “AUGUAG,” or “AUGUGA,” with a single methionine. For example, the NCYM transcript has a pPTI with a length of 109 in frame 1, three sPTIs with lengths of 69, 8, and 6, respectively, in frame 2, and no PTIs in frame 3 (Supplementary Figure 1C and D).

#### Characteristics

Therefore, the following relationship between the lengths of pPTI and sPTI is held:

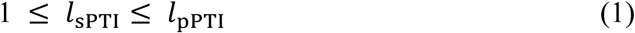

### PTI score

#### Definition

We defined the PTI score (Figure 1A) according to Equations 2 and 3.

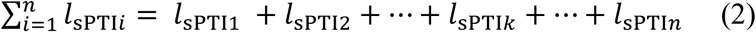

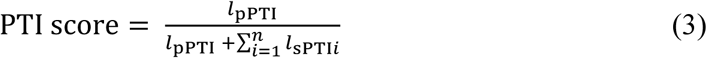

where 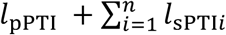 represents the sum of all PTI lengths. The definition is derived from the hypothesis that the potential for translation of a pPTI is reduced by translation of sPTIs. Consistent with this hypothesis, coding transcripts with translation of sPTIs had lower PTI scores than all coding transcripts (Figure 2C).

#### Example

For an RNA sequence with only one PTI, the PTI score is 1 (Figure 1B). An RNA sequence with many sPTIs tended to have a score close to 0 (Figure 1B). If the sum of all sPTI lengths was equal to pPTI length, the PTI score is 0.5 (Figure 1B). The PTI score of the *NCYM* transcript is 0.568 (Supplementary Figure 1C). Further information is included in the Supplementary Notes.

#### Characteristics

Therefore, the range of the PTI score is:

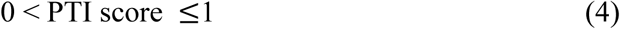

### Relative frequencies *f*(*x*) and *g*(*x*)

#### Definition

We defined the, *f*(*x*) and *g*(*x*), respectively, as (Figure 1C):

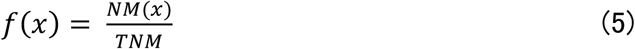

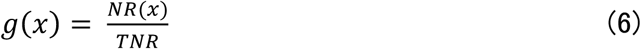

where *TNM* and *TNR* represent the total numbers of coding and noncoding transcripts, respectively, excluding transcripts lacking PTIs. *NM*(*x*) and *NR*(*x*) are the numbers of coding and noncoding transcripts with a PTI score of *x*, respectively. To define coding/non-coding transcripts with a PTI score of *x*, we divided the histograms into ten classes, and used the median values of the classes to represent the PTI score (Figure 1C). Therefore, in Equations 5 and 6, the PTI score *x* is restricted as follows:

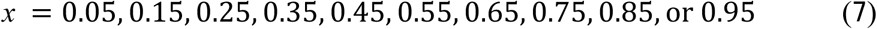

#### Characteristics

Thus, *f*(*x*) and *g*(*x*) follow Equations 8–11:

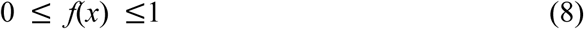

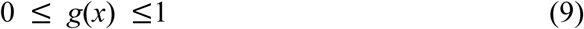

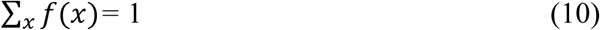

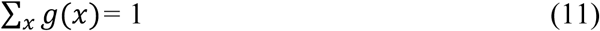

### Overlapping scores *O*_pti_ and *O*_cov_

#### Definition

The *O*(*x*) was calculated according to Equation 12:

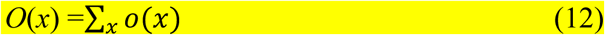

where *o*(*x*) is the smaller value of the relative frequency of coding *f*(*x*) or noncoding transcripts *g*(*x*). *O*_pti_ is *O*(*x*) with PTI score = *x,* and *O*_cov_ is *O*(*x*) with ORF coverage = *x*.

### Protein-coding potential *F*(*x*)

#### Definition

*F*(*x*) was calculated according to Equation 13:

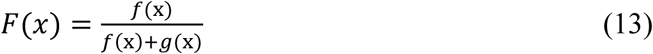

#### Example

For example, *F*(0.15) in human transcripts is shown in Figure 1D. *F*(0.15) was calculated using Equation 13, as follows:

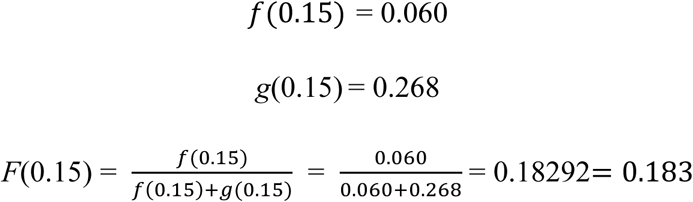

### Identification of noncoding transcripts with high protein-coding potential

NR transcripts with high *F*(*x*) (0.6 ≤ *x* < 0.8) were identified from the total NR transcripts from the NCBI nucleotide database. NR transcripts shorter than 200 nucleotides or with pPTIs encoding putative peptides with fewer than 20 residues were excluded. The amino acid sequences of pPTIs in these transcripts were subjected to a BLASTP search to detect the presence of putative domain structures. In the BLASTP search, non-redundant protein sequences (nr) were applied as the search set, and quick accelerated protein-protein BLAST (BLASTP) was chosen as the algorithm. In the search results, putative conserved domains or the message “No putative conserved domains have been detected” are shown in the Graphical Summary tab.

CDSEARCH/cdd was used to search for conserved domain structures using the default settings: low-complexity filter, no; composition-based adjustment, yes; E-value threshold, 0.01; maximum number of hits, 500. Based on these data, transcripts with or without putative conserved domain structures are indicated as + or -, respectively.

### Functional annotation of original genes

Original genes were defined as those noted in the official gene name of NR transcripts, including sense genes for antisense transcripts, homologous genes for pseudogenes, coding genes for noncoding transcript variants, and readthrough, divergent, or intronic transcripts. For lincRNAs, miRNA host genes, small nuclear RNAs, and other lncRNAs, the official gene symbol was used for annotation. This information was manually checked using the information available in the nucleotide database. The DAVID program (https://www.david.ncifcrf.gov) was used to identify the enriched molecular functions and pathways related to the original genes. *Q*-values (*P*-values adjusted for false discovery rate) were calculated using the Benjamini–Hochberg method in DAVID.

### Nonsynonymous (*K*a) to synonymous (*K*s) nucleotide substitution ratios

To identify orthologous regions between human transcripts and chimpanzee/mouse genomes, BLAT v. 36 (Kent 2002) was conducted using human transcript sequences with the estimated PTI score against chimpanzee (PtRV2) and mouse (GRCm38.p6) genomic sequences defined in the NCBI database. We defined the blat best-hit genomic regions of chimpanzee/mouse as orthologs for each human transcript. The human– chimpanzee (or human–mouse) sequences were aligned for each exon region and the sequences were combined for each transcript. Only orthologous sequence pairs of more than 60 bp in length (encoding > 20 amino acid residues) were extracted.

Nonsynonymous (*K*a) and synonymous (*K*s) nucleotide substitution rates were estimated as described by Yang and Nielsen (Yang and Nielsen 2000), implemented in PAML version 4.8a (Yang 1997). Transcripts with high *K*a (> 1) or high *K*s (> 1) were excluded from our dataset as outliers. We calculated *K*a and *K*s for 47,228 NM human– chimpanzee, 14,116 NM human–mouse, 8,810 NR human–chimpanzee, and 1,561 NR human–mouse pairs.

### Relative frequencies of negatively selected genes

We defined this frequency, *h*(*x*), in coding and noncoding transcripts (Figure 1G), as shown in Equation 14:

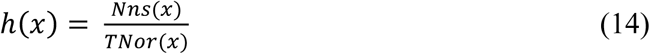

where *TNor*(*x*) represents the total number of coding or noncoding transcripts with orthologous sequences at PTI score = *x*. *Nns*(*x*) is the number of coding or noncoding transcripts with *K*a/*K*s < 0.5 at PTI score = *x*. The PTI score *x* is restricted as shown in Equation 7.

### Phylogenic trees

TimeTree (Hedges et al. 2006) was used to draw trees using official species names.

### Selection of viruses and identification of vORFs

The complete genomes of positive-sense single-stranded RNA viruses infecting humans or bacteria (Supplementary Tables 14 and 15) were collected from the NCBI Virus database (Hatcher et al. 2017). Viral ORFs were identified, and the sums of vORF lengths 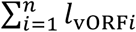 were manually calculated. We eliminated those viruses that translated viral proteins after splicing or using exceptional translation mechanisms such as ribosome frameshifting, alternative initiation sites, ribosome slippage, and RNA editing.

### vORF score

#### Definition

The vORF score was calculated according to Equations 15–17:

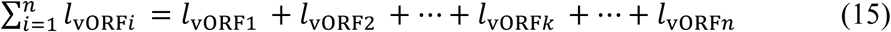

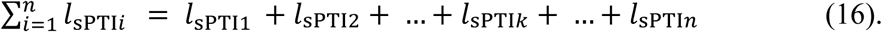

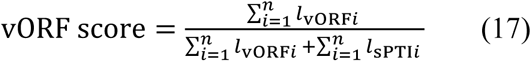

where *l*_vORF*i*_ represents the length of the bona fide ORFs, and 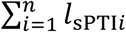 is the sum of the lengths of the secondary PTI lengths. 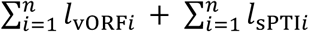 represents the sum of the lengths of all PTIs, including all ORFs.

### PTI score calculations using transcriptome data

Transcriptome data from five species were obtained from a previous study (Sarropoulos et al. 2019). All transcripts expressed at detectable levels (non-zero) in each tissue were used to calculate PTI scores for lncRNAs and to plot PTI score distributions. For the correlation between tissue specificity and PTI score, we divided the transcripts into the indicated groups according to the number of tissues in which the transcript was detected and described the PTI score distribution in each group. Human transcriptome data for coding transcripts were obtained from the Human Protein Atlas (http://www.proteinatlas.org), including RNA isoform data from 131 cell lines and 281 tissues. The PTI score for each transcript was calculated from Ensembl data.

### Statistical analyses

Statistical analyses were performed using Excel and R software (R Project for Statistical Computing, Vienna, Austria).

### Data availability

1. Source data for statistical analyses and figures (10 example datasets): https://figshare.com/s/498cb340a075284b2dbf
2. Code associated with generating and analyzing these tables: https://figshare.com/s/0f1ed0954d5bd620eb59

## Acknowledgments

We thank Yohko Yamaguchi, Yoshitaka Hippo, Sana Yokoi, and Akira Nakagawara for their helpful comments; Isao Kurosaka and Miayamoto Joe Philips for assistance in calculating PTI scores, and Mayuko Seto, Ryo Ohtaka, Manami Shinbori, Abedin Swapnil Eishraq, Kanna Kamikawa, and Taichi Yokoi for assistance in checking putative domain structures using BLASTP and classification of species according to the IUCN Red List. This work was partially supported by the Yasuda Medical Foundation, Takeda Science Foundation, and Interstellar Initiative grant18jm0610006h0001 to Y.S. from the Japan Agency for Medical Research and Development.

## Author contributions

Y.S. conceived and developed the research plan; Y.S., M.K., M.N., K.N., H.K., M.K., and T.M. analyzed the data; and Y.S., M.K., and T.M. wrote the manuscript.

## Additional information

Supplementary information is available.

## Supplementary figure legends

**Supplementary Figure 1. PTIs of *NCYM* and an example of PTI score calculation.** (A) *NCYM* cDNA sequence. (B) Coding prediction of *NCYM* by CPAT (http://lilab.research.bcm.edu). (C) Translated amino-acid sequence of *NCYM* in 3 frames in the 5ʹ to 3ʹ (sense) direction. Red characters, primary PTI; blue characters, secondary PTIs. Stop codons are shown as asterisks. (D) Calculation of PTI score and *F*(*x*) for the *NCYM* transcripts. The length of pPTI is 109 and the sum of sPTI lengths is 83; therefore, the PTI score is 0.568.

**Supplementary Figure 2. Relationships between relative frequencies of coding and noncoding transcripts for human and mouse PTI scores.** (A) Histogram of PTI score relative frequencies in coding *f*(*x*) and noncoding *g*(*x*) human transcripts with random shuffling controls using human data sets from RefSeq. (B) Relative frequencies of coding *f*(*x*) and noncoding *g*(*x*) transcripts calculated using human data sets from Ensembl. (B) Relative frequencies of coding *f*(*x*) and noncoding *g*(*x*) transcripts calculated using mouse data sets from RefSeq (upper panels) or Ensembl (lower panels).

**Supplementary Figure 3. PTI scores correlate with protein-coding potential, *F*(*x*), at PTI scores** ≤ **0.65 for human and mouse transcripts.** (A) Relationship between PTI score and *F*(*x*) in a human data set from Ensemble and random controls (center). Random shuffling controls (right) were generated from a human data set from both Ensemble and RefSeq. (B) Relationships between PTI score and *F*(*x*) in mouse transcripts using data sets from RefSeq (upper panels) or Ensembl (lower panels).

**Supplementary Figure 4. Effects of translation on distributions of ORF coverage and size in human lincRNAs.** ORF coverage (A) and ORF size (B) distributions of lincRNAs encoding small proteins (red line, n = 174) registered in the SmProt database (http://bioinfo.ibp.ac.cn/SmProt/) compared with all lincRNAs registered in Ensembl (black line, n = 11,875).

**Supplementary Figure 5. Translation of sPTI affects on human PTI score distributions.** PTI score distributions of coding transcripts with translation of uORFs (blue, n = 14,506), sORFs (red, n = 80), and dORFs (green, n = 3,955) registered in the sORF database (http://www.sorfs.org/), compared to all coding transcripts registered in Ensembl (black, n = 94,039).

**Supplementary Figure 6. Relationships between PTI scores and relative frequencies of coding and noncoding transcripts in archaea**. Phylogenetic tree for 9 archaeal species and histogram of *f*(*x*) (white) or *g*(*x*) (black) in the data (left) and in nucleic-acid–scrambled controls (right). PTI scores with highest *f*(*x*) are indicated in the histograms. The lineage of one archaea species (*Nitrososphaera viennensis* EN76) is unknown and thus excluded from the phylogenic tree. *O*_pti_ was calculated using the PTI score distribution of observed data.

**Supplementary Figure 7. Relationships between PTI score and relative frequencies for coding and noncoding transcripts in plants.** Phylogenetic tree for 12 plants and histogram of *f*(*x*) (white) or *g*(*x*) (black) in the data (left) and in sequence–scrambled controls (right). PTI scores with highest *f*(*x*) are indicated. *O*_pti_ was calculated using the PTI score distribution of observed data.

**Supplementary Figure 8**. **Relationships between PTI score and relative frequencies of coding and noncoding transcripts in species shown in** Figure 6. Phylogenetic tree for 24 cellular organisms and histograms of *f*(*x*) (white) or *g*(*x*) (black) for PTI scores (left) and ORF coverage (right). *O*_pti_ and *O*_cov_ were calculated using the distribution of observed data.

**Supplementary Figure 9**. **Relationship between overlaps of ORF coverage distributions and effective population sizes.** Dot plots of *O*_cov_ and *U*_p_ (left) and *N*_e_ (right). These relationships are approximated by the logaritic and exponential functions, respectively. White, gray, and black dots indicate bacteria, unicellular eukaryotes, and multicellular eukaryotes, respectively.

**Supplementary Figure 10. Relationships between PTI scores and protein-coding potential *F*(*x*) for 32 eukaryotes.** Data sets from Ensembl (left, used in Figure 7) and random controls (right). Mouse and human data are identical to those shown in Supplementary Figure 3. Shapes of approximate functions are shown as L or C, indicating linear (in black) and constant (in red) functions, respectively. Numbers of lncRNAs with PTI score 0.05 were < 5 in *U. americanus*, *C. canadensis*, and *G. gorilla*. Therefore, we eliminated the *F*(0.05) from these species for the approximation by linear functions (asterisks).

**Supplementary Figure 11. PTI score distributions of lncRNAs from tissues from four mammals.** PTI score distributions for mature testes and other tissues are indicated as black and gray lines, respectively.

**Supplementary Figure 12. Relationships between tissue specificity and PTI score distributions for noncoding transcripts from four mammals.** Line intensity represents specificity of gene expression.

**Supplementary Figure 13. Relationships between tissue-specificity and PTI score distributions for coding transcripts from human cell lines (A) or tissues (B).** Line intensity represents specificity of gene expression.

**Supplementary Table 1. Studied organisms with official names, taxonomic IDs, lineage information, and numbers of coding/noncoding transcripts.**

**Supplementary Table 2. Human NR transcripts reassigned as NM during the past three years.**

**Supplementary Table 3. Human NR transcripts with high protein-coding potential (0.6** ≤ **PTI score < 0.8).**

**Supplementary Table 4. Functional annotation of human NR transcripts with high protein-coding potential and without putative domain structure(s).**

**Supplementary Table 5. Functional annotation of human NR transcripts with high protein-coding potential and with putative protein domain structure(s).**

**Supplementary Table 6. Mouse NR transcripts with high protein-coding potential (0.6** ≤ **PTI score < 0.8).**

**Supplementary Table 7. Functional annotation of mouse NR transcripts with high protein-coding potential and without putative protein domain structure(s).**

**Supplementary Table 8. Functional annotation of mouse NR transcripts with high protein-coding potential and with putative domain structure(s).**

**Supplementary Table 9. NR transcripts with high protein-coding potential (0.6** ≤ **PTI score < 0.8) from *C*. *elegans*.**

**Supplementary Table 10. Functional annotation of NR transcripts with high protein-coding potential and without putative domain structure(s) from *C*. *elegans*.**

**Supplementary Table 11. Functional annotation of NR transcripts with high protein-coding potential and with putative domain structure(s) from *C*. *elegans*.**

**Supplementary Table 12. Twenty-three human noncoding transcripts showing both negative selection (*K*a/*K*s < 0.5) and high PTI scores.**

**Supplementary Table 13. Organisms shown in** Figure 6 **with official names, taxonomy IDs, lineage information, and numbers of coding or noncoding transcripts, *O*_pti_, and *O*_cov_. The effective population sizes (*N*e) and mutation rates (*U*p) were estimated by Lynch et al. (2016).**

**Supplementary Table 14. Positive-sense, single-stranded human viruses with official names, taxonomy IDs, lineage information, genome lengths, and sequences.**

**Supplementary Table 15. Positive-sense, single-stranded bacteriophages with official names, taxonomy IDs, lineage information, source information, genome lengths, and sequences.**

