## Supplementary Figures for "Protein-coding potential of RNAs measured by potentially translated island scores"

### Supplementary figure 1

A

```
>NR_161162.1 Homo sapiens MYCN opposite strand (MYCNOS),
transcript variant 2, long non-coding RNA
CATTTCATTCACACAAGGCACTGCCTGGGGGAGGGGGCTGTTCCTGGCTGCAGAATTCTAGCTCTCACGA
GCACGCAGACAACCGCACTCGCAGCGGTGTGGGGCCGGCTGCTCAGGGGAAGCCCCAGGCTCTCCGACCC
AGCTACCGGGAATGGGGCACCCTTTGGAGAAGAACCCCAGCCTGGGGTGGGGACGCACCGGCTCTCCGAC
AGCTCAAACACAGACAGATCTTCTAGAGCCGAGGGAATTTCTTTTCGCAGAAGCCATTACTCCCCCCGAG
AGAAGGCTGCAAAGCTGGGAAGCCCAGGGTGTGCTCCTCCCGCCCTTTTGGACCCCCGGGCTTGCACCGG
CTGCACTCTGAGAACCAGCTGCGCGCGGAGCGGTGCAATGCAGCACCCACCCTGCGAGCCTGGCAATTGC
TTGTCATTAAAAGAAAAAAAAATTACGGAGGGCTCCGGGGGTGTGTGTTGGGGAGGGGAGACCGATGCTT
CTAACCCAGCCCCCGCTTTGACTGCGTGTTGTGCAGCTGAGCGCGAGGCCAACGTTGAGCAAGGCCTTGC
AGGGAGGTTGCTCCTGTGTAATTACGAAAGAAGGCTAGTCCGAAGGTGCAAAATAGCAGGGAGAGGACGC
GCCCCCTTAGGAACAAGACCTCTGGATGTTTCCAGTTTCAAATTGAAAGAAGAGGGGCGCCCCCCTTGTT
TGAAAATAAATAAATAAATAAGTGCGAGCTAC
```

B

| Sequence Name | ORF size | Ficket Score | Hexamer Score | Coding Probability | Coding label |
|---|---|---|---|---|---|
| NR_161162.1 | 330 | 0.504 | -0.1207753 | 0.022153 | no |

C

5'3' Frame 1

HFIHTRHCLGEGAVPGCRILALTSTQTTALAAVWGRLLRGSPRLSDPATGNGAPFGEEPQPGV
GTHRLSDSSNTDRSSRAEGISFRRSHYSPREKAAKLGSPGCAPPALLDPRACTGCTLRTSCAR
SGA**MQHPPCEPGNCLSLKEKKITEGSGGVCWGGETDASNPAPALTACCAAEREANVEQGLAGR**
**LLLCNYERRLVRRCKIAGRGRAPLGTRPLDVSSFKLKEEGRPPCLKINK ***ISASY

5'3' Frame 2

ISFTQGTAWGRGLFLAAEF*LSRARRQPHSQRCGAGCSGEAPGSPTQLPG**MGHPLEKNPSLGW**
**GRTGSPTAQTQTDLLEPREFLFAEAITPPERRLQSWEAQGVLLPPFWTPGLAPAAL***EPAARG
AVQCSTHPASLAIACH*KKKKLRRAPGVCVGEGRP**MLLTQPPL***LRVVQLSARPTLSKALQGG
CSCVITKEG*SEGAK*QGEDAPP*EQDLW**MFPVSN***KKRGAPLV*K*INK*VRA

5'3' Frame 3

FHSHKALPGGGGCSWLQNSSSHEHADNRTRSGVGPAAQGKPQALRPSYREWGTLWRRTPAWGG
DAPALRQLKHRQIF*SRGNFFSQKPLLPPREGCKAGKPRVCSSRPFGPPGLHRLHSENQLRAE
RCNAAPTLRAWQLLVIKRKKNYGGLRGCVLGRGDRCF*PSPRFDCVLCS*ARGQR*ARPCREV
APV*LRKKASPKVQNSRERTRPLRNKTSGCFQFQIERRGAPPLFENK*INKCEL

D

$$l_{pPTI} = 109$$

$$\sum_1^3 l_{\mathrm{sPTIk}} = 69 + 8 + 6 = 83$$

$$\text{PTI score} = \frac{109}{109+83} = 0.568$$

$$F(x) = 1.301 \times 0.568 + 0.0072 = 0.746 \text{ (Ensemble)}$$

$$F(x) = 1.313 \times 0.568 + 0.0189 = 0.765 \text{ (RefSeq)}$$

### Supplementary figure 2

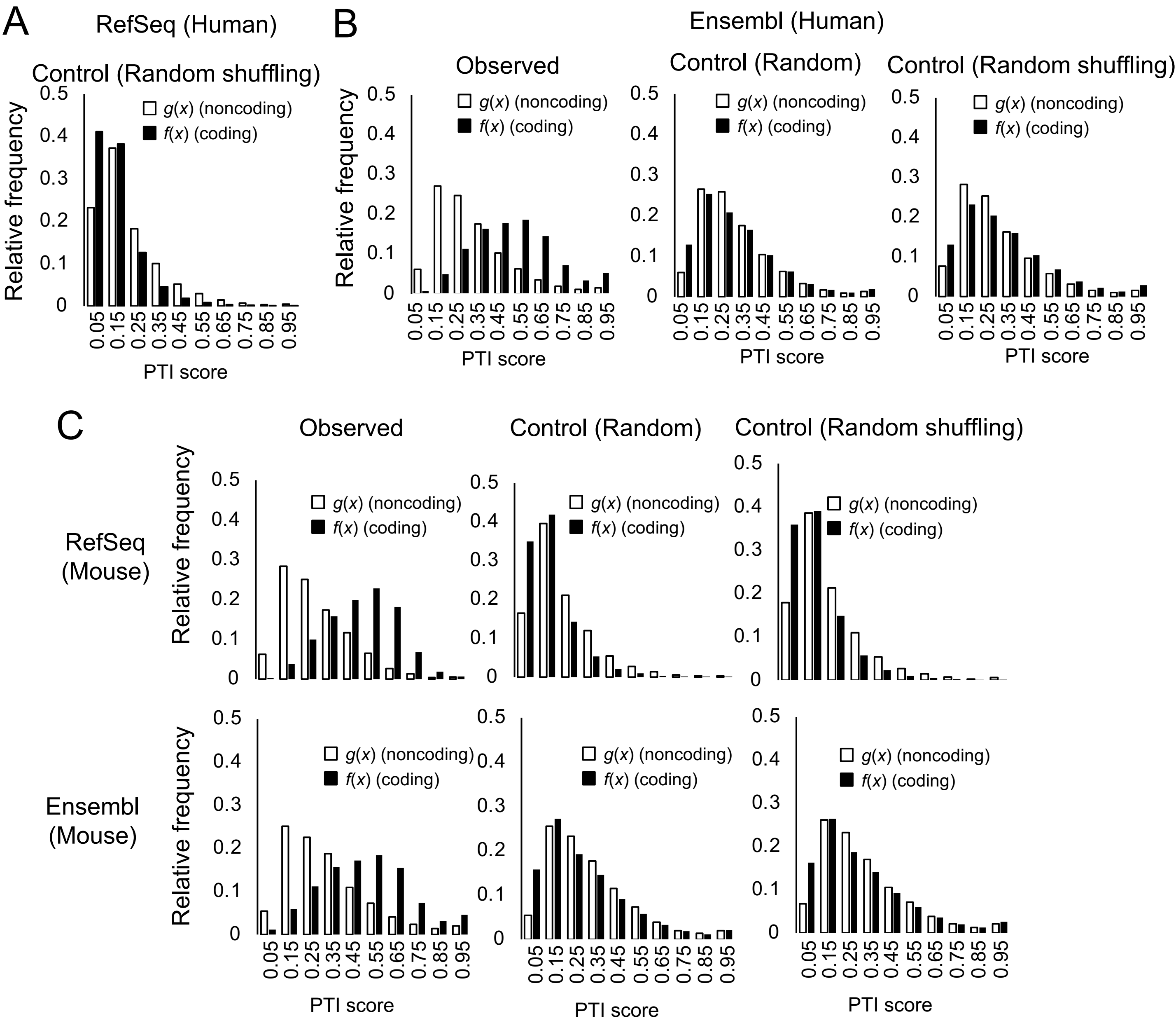

**Supplementary figure 3**

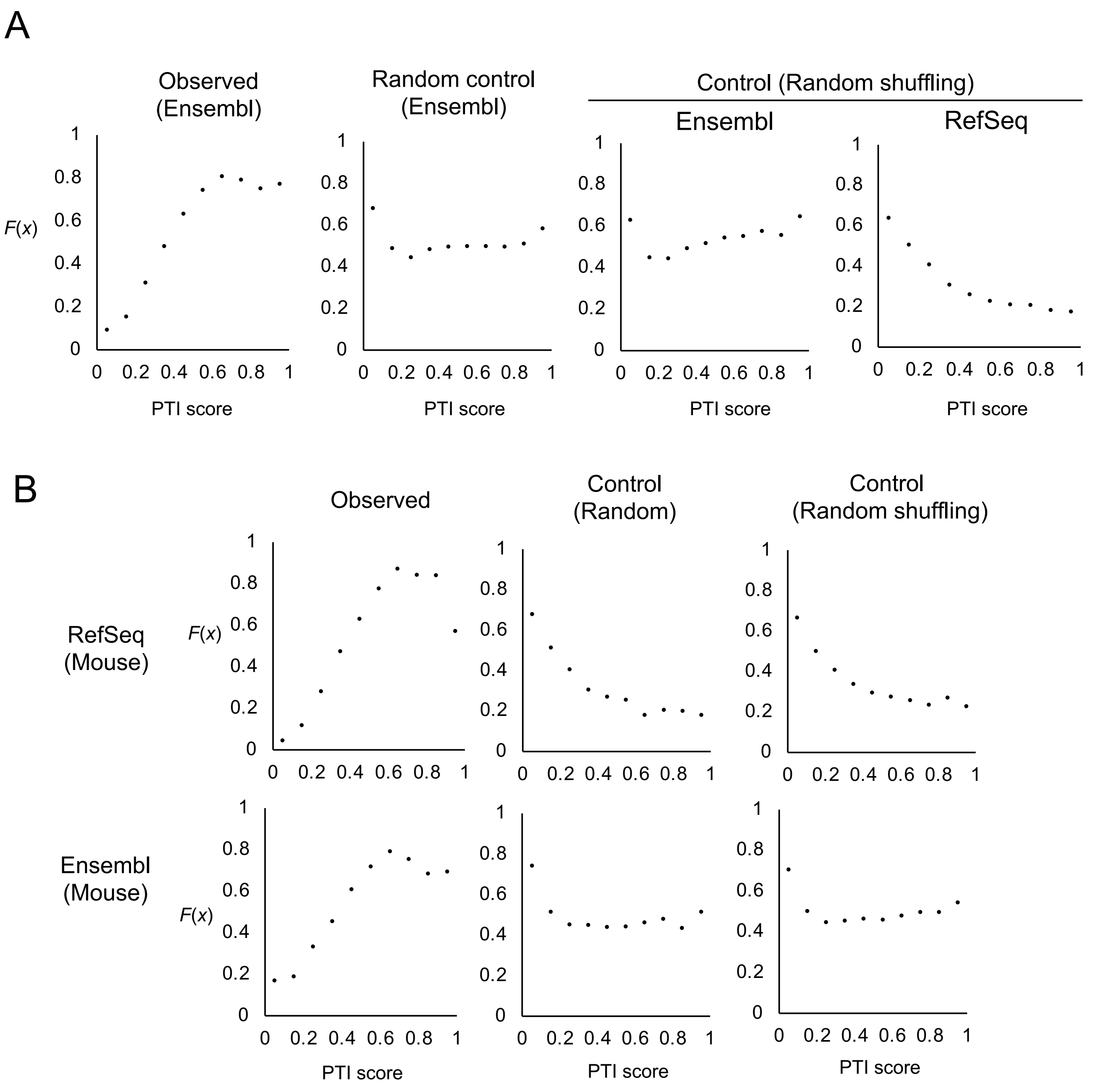

#### Supplementary figure 4

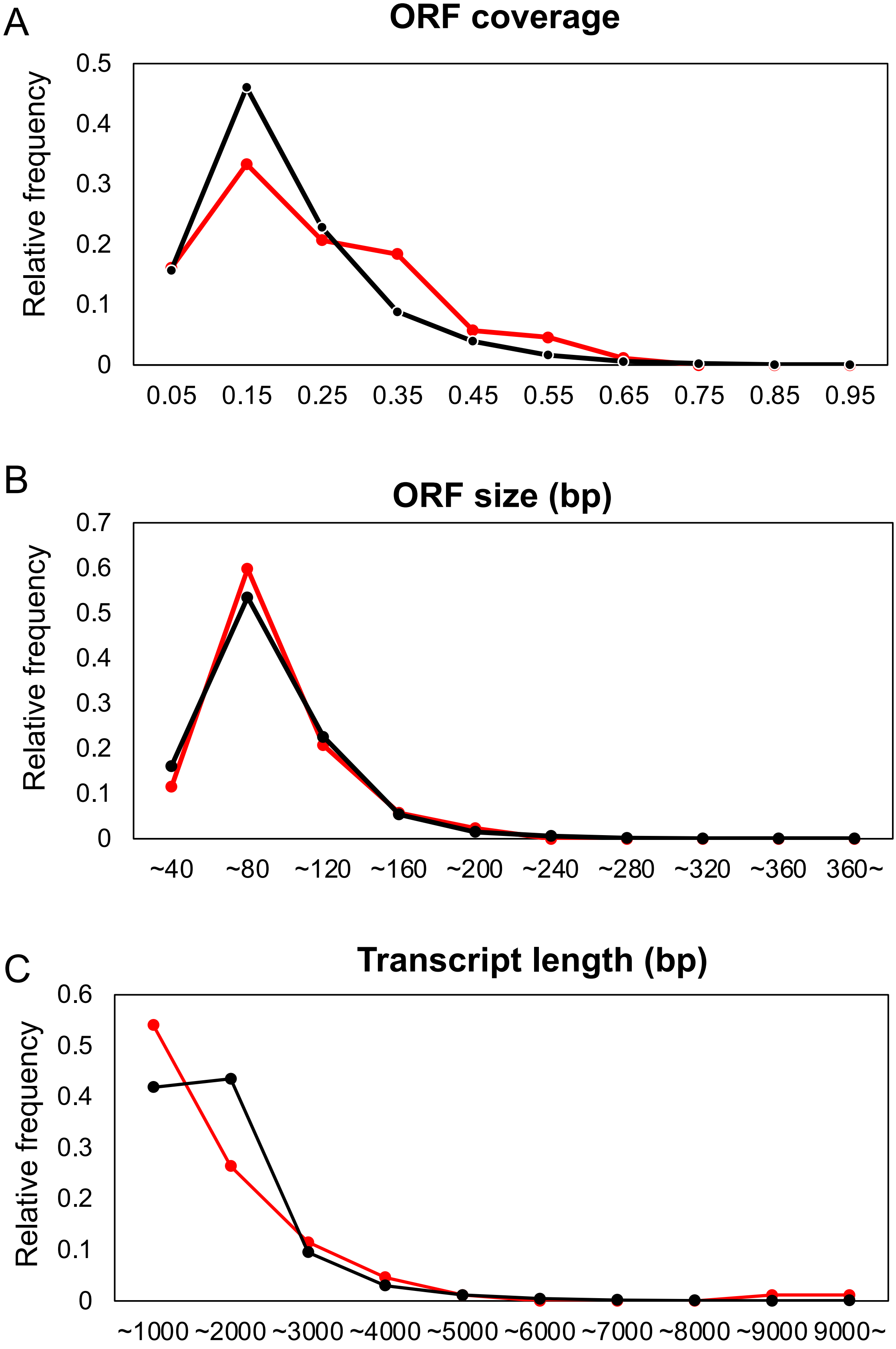

### Supplementary figure 5

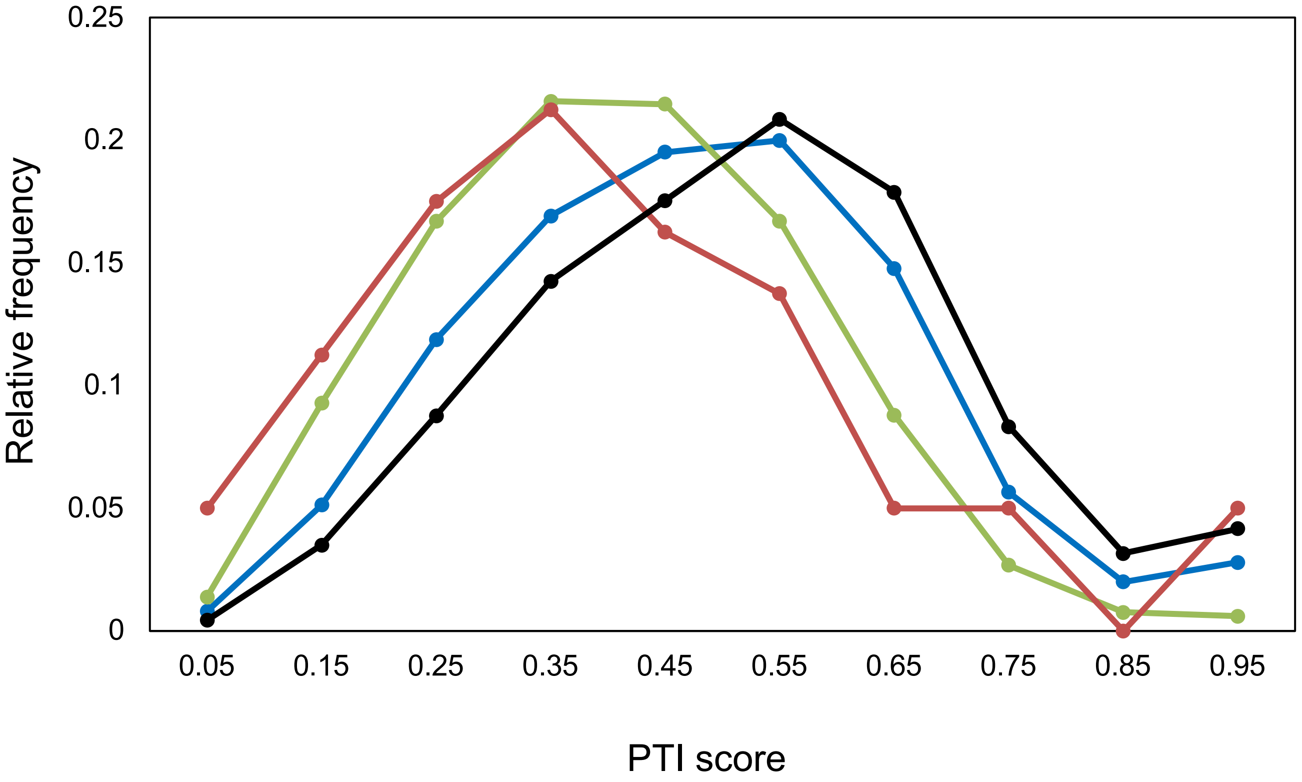

### Supplementary figure 6

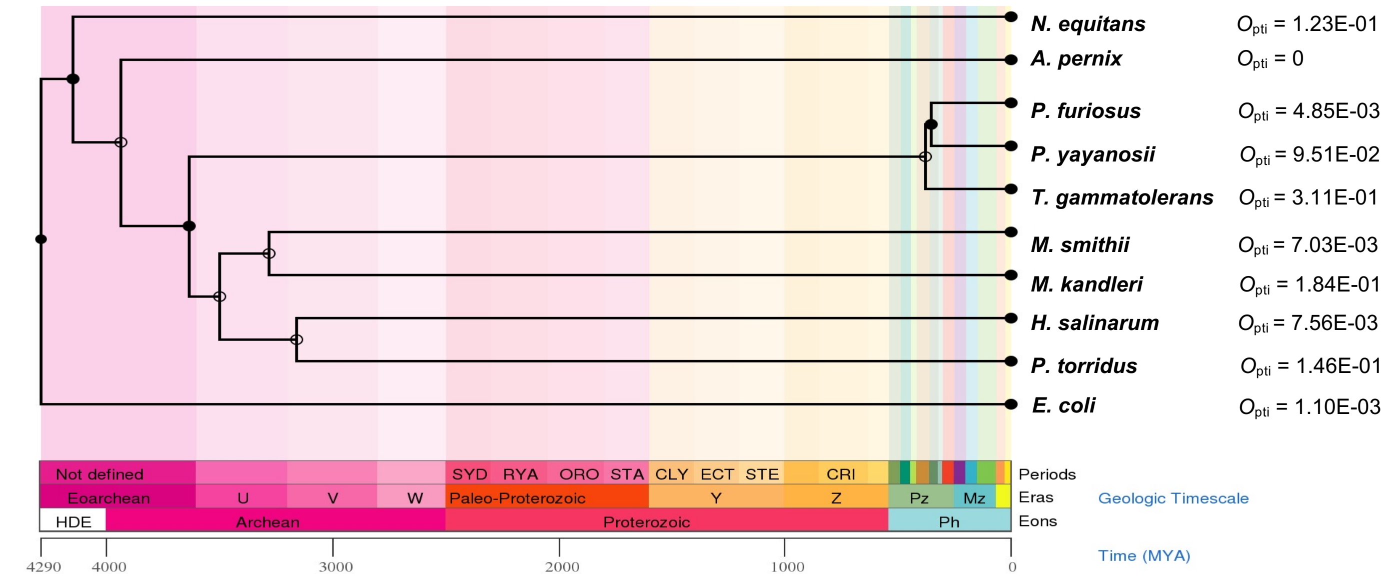


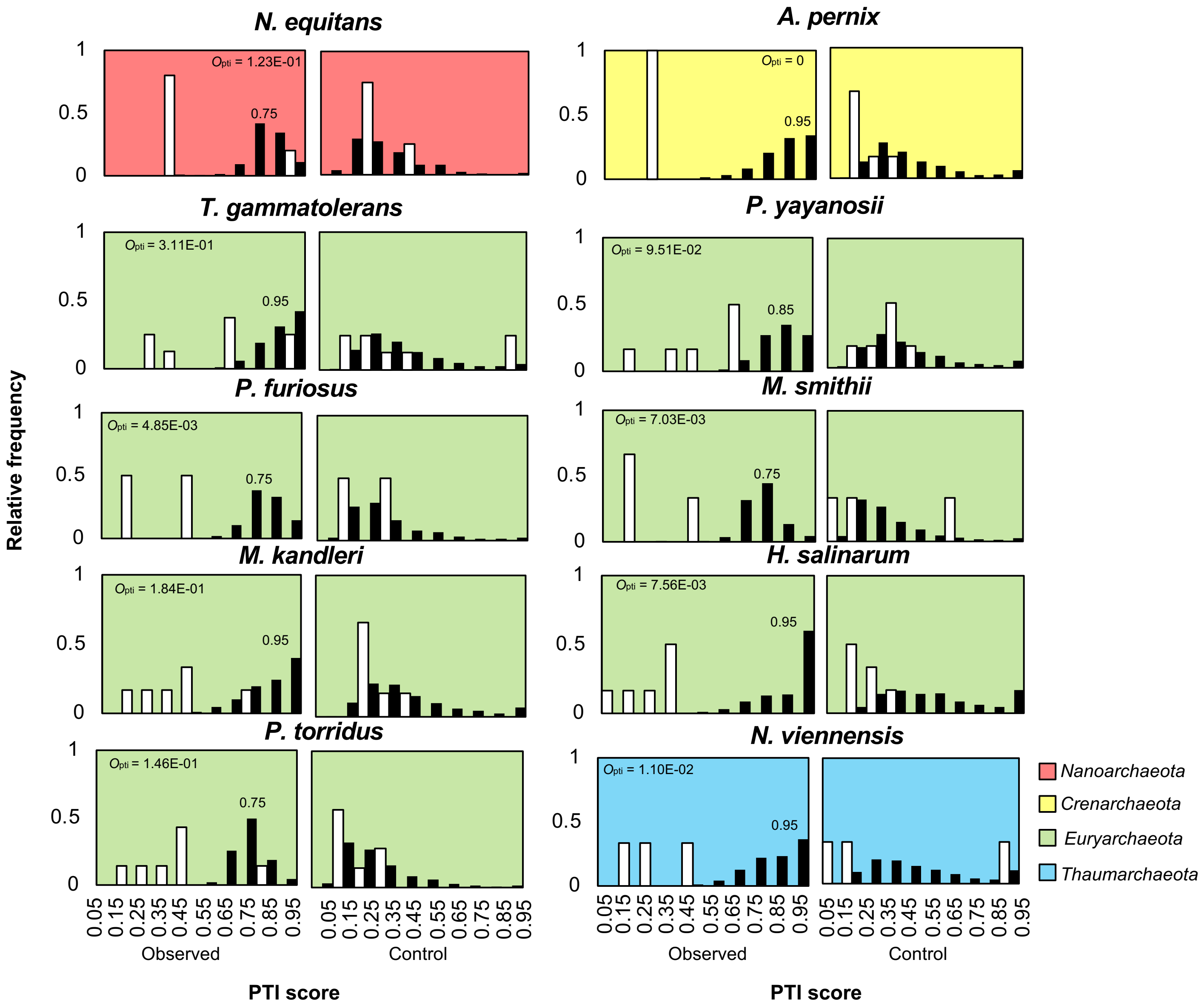

### Supplementary figure 7

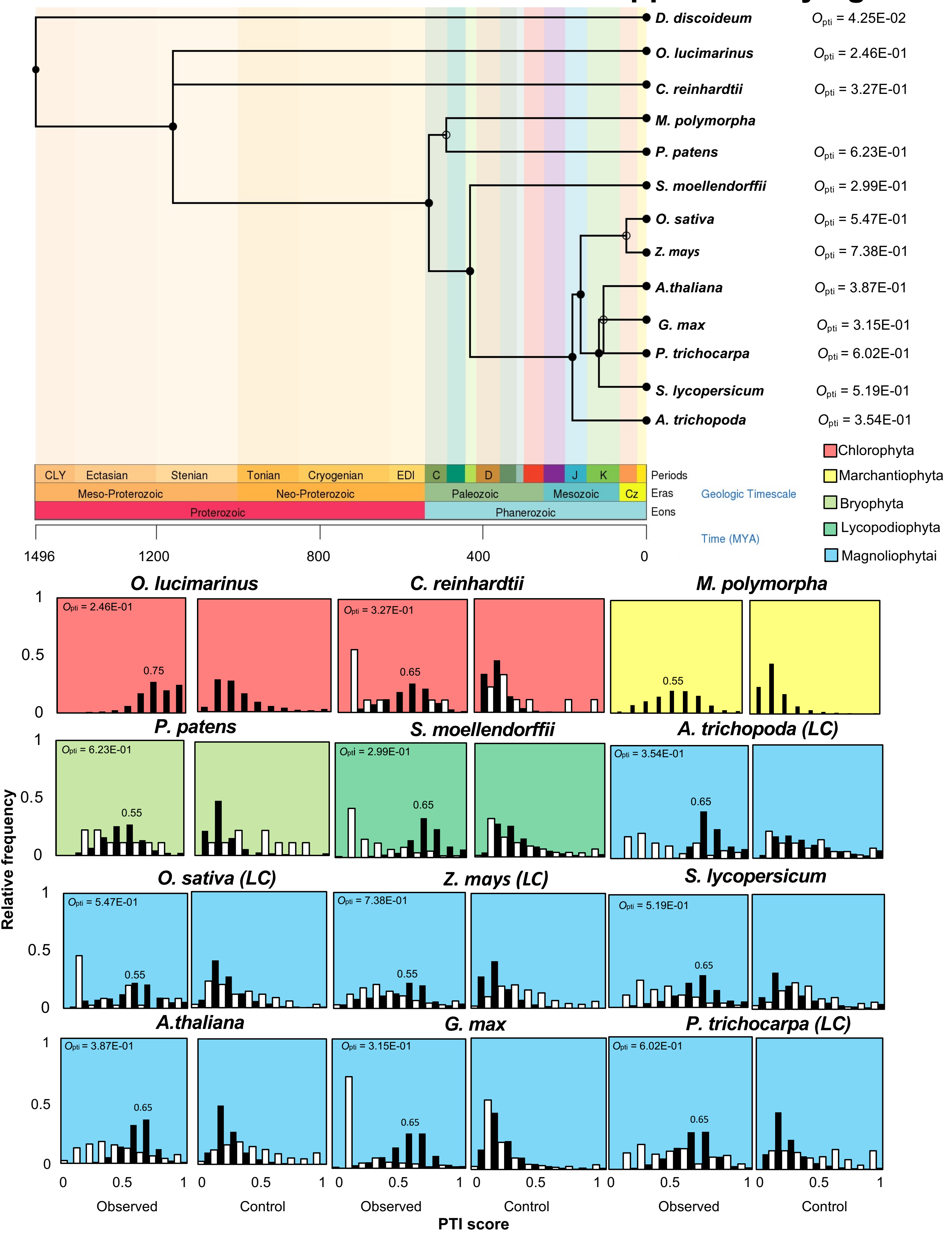

### Supplementary figure 8

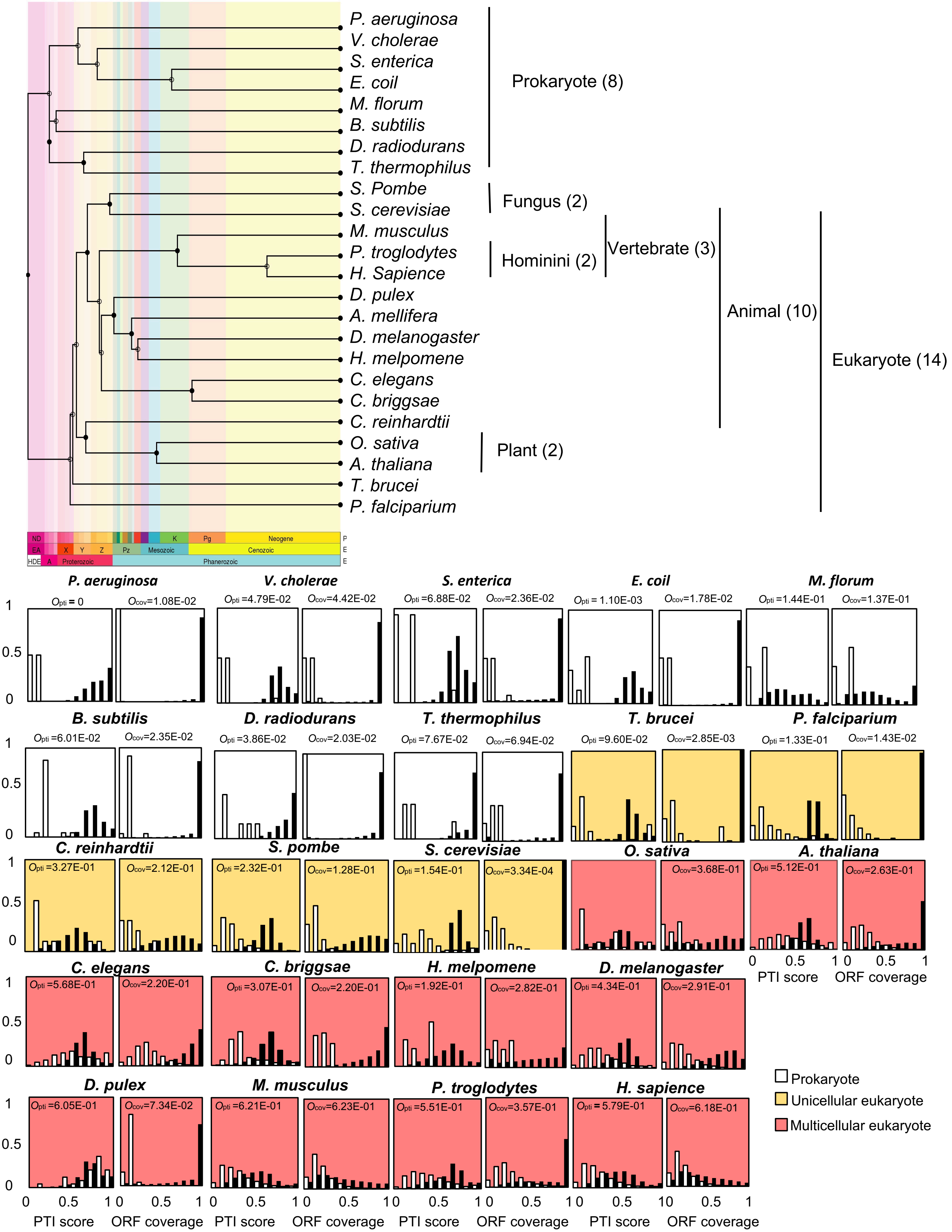

### Supplementary figure 9

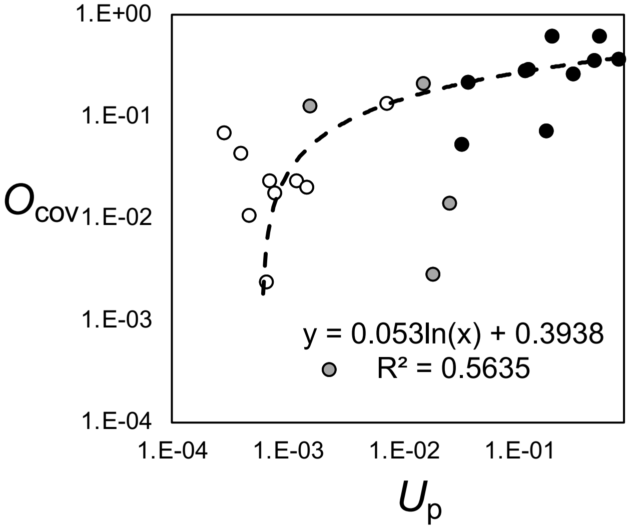


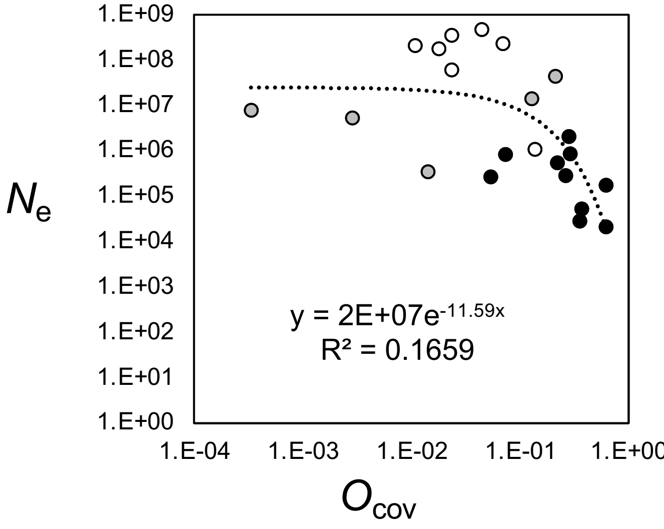

### Supplementary figure 10

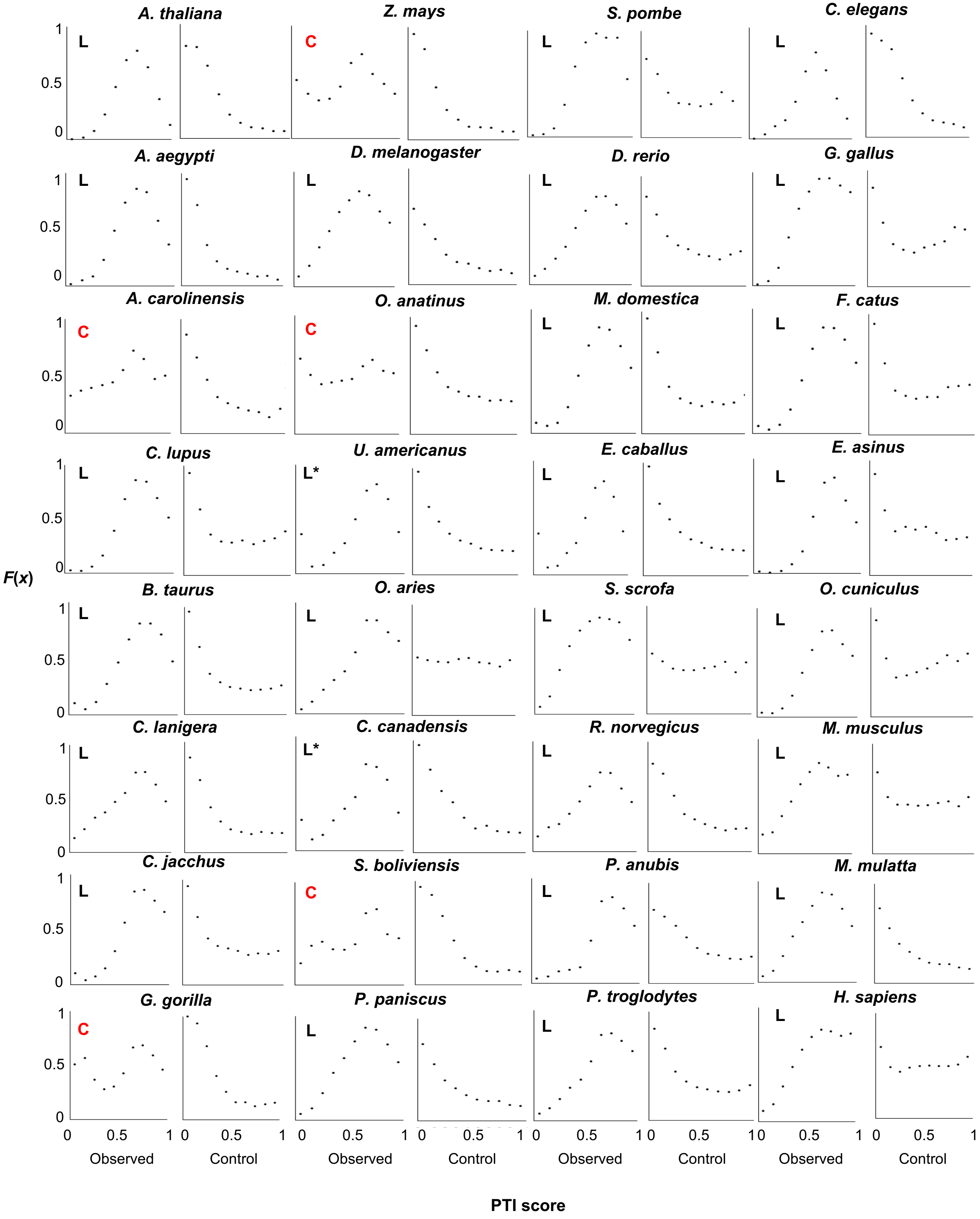

### Supplementary figure 11

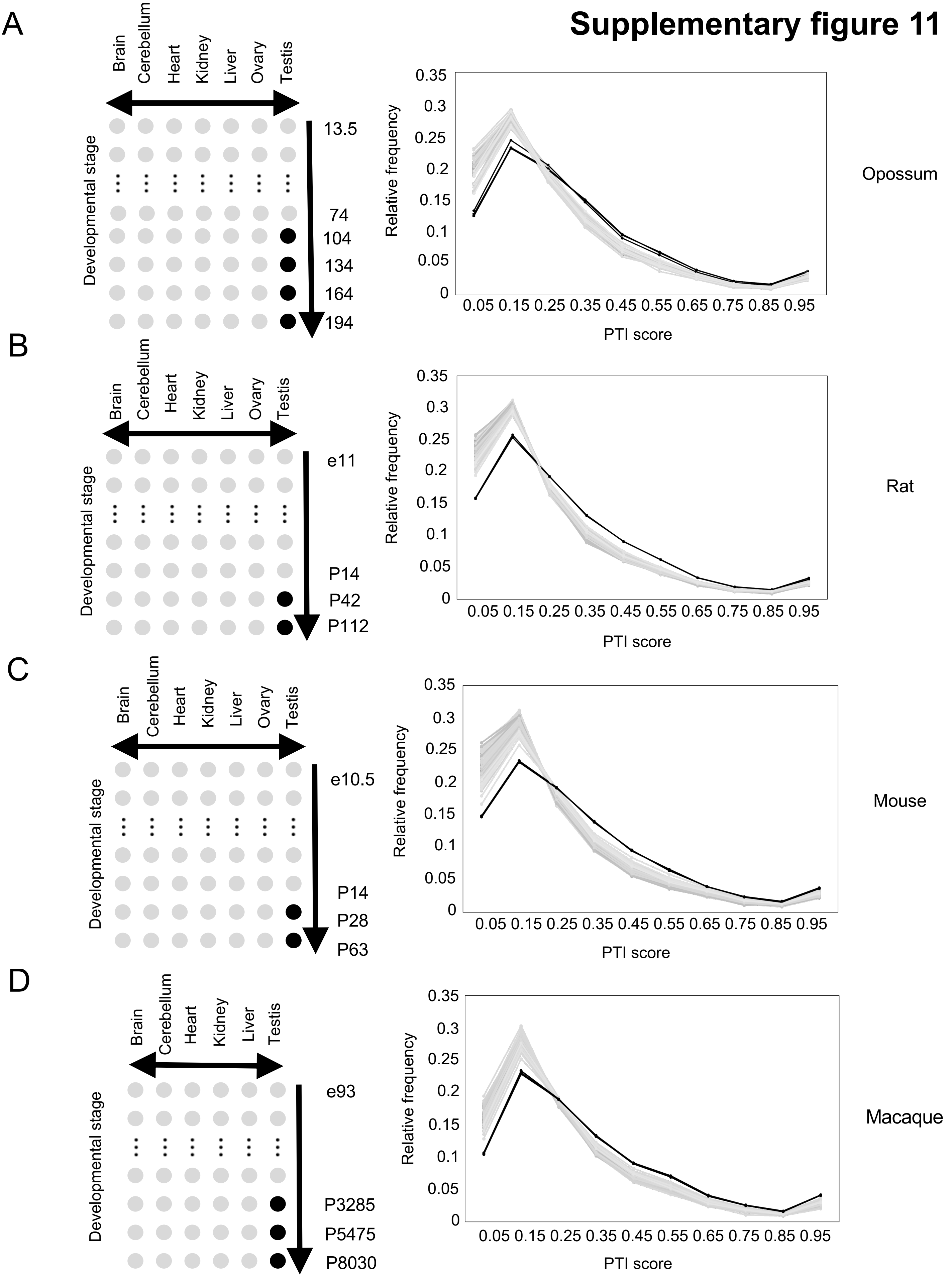

### Supplementary figure 12

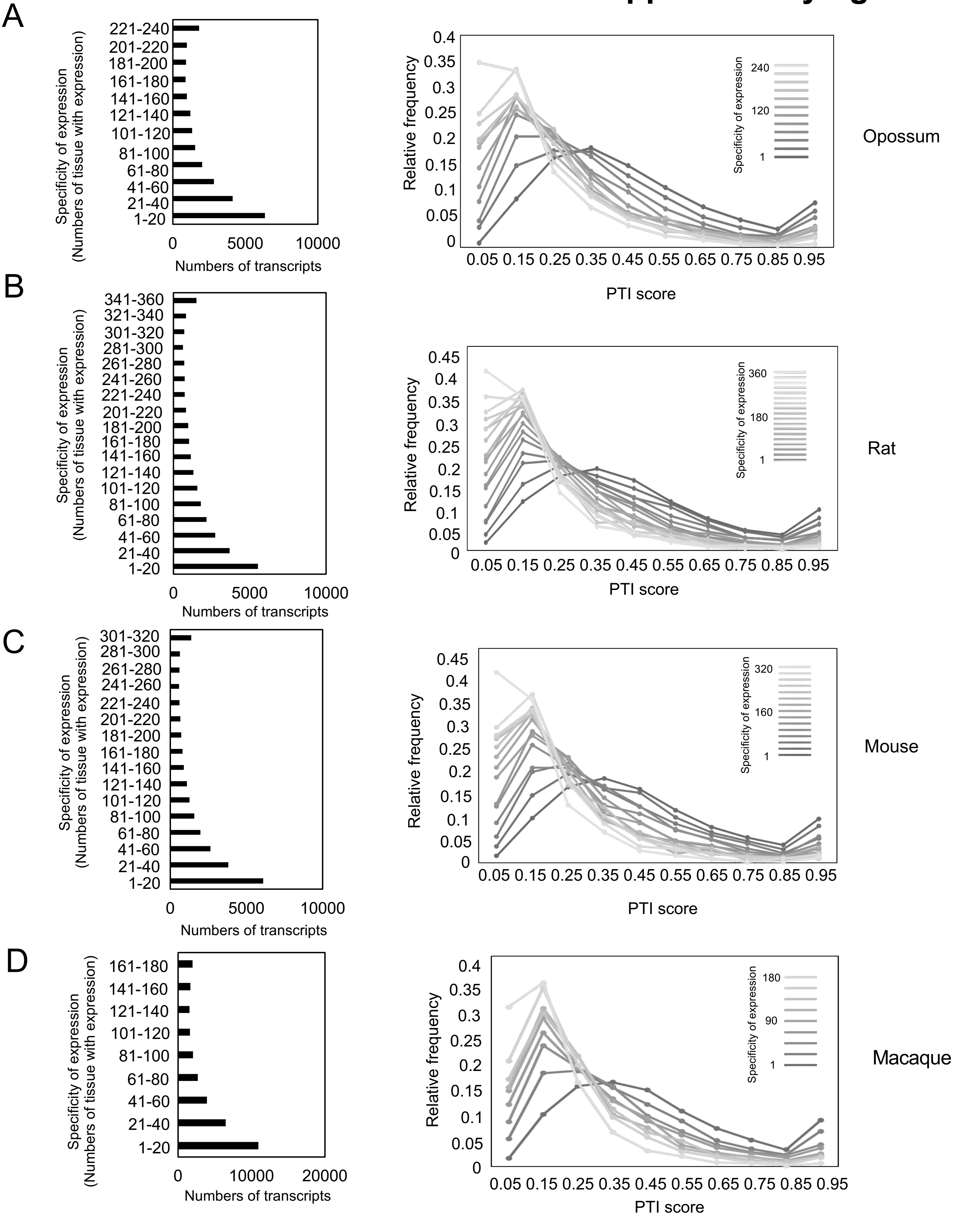

### Supplementary figure 13

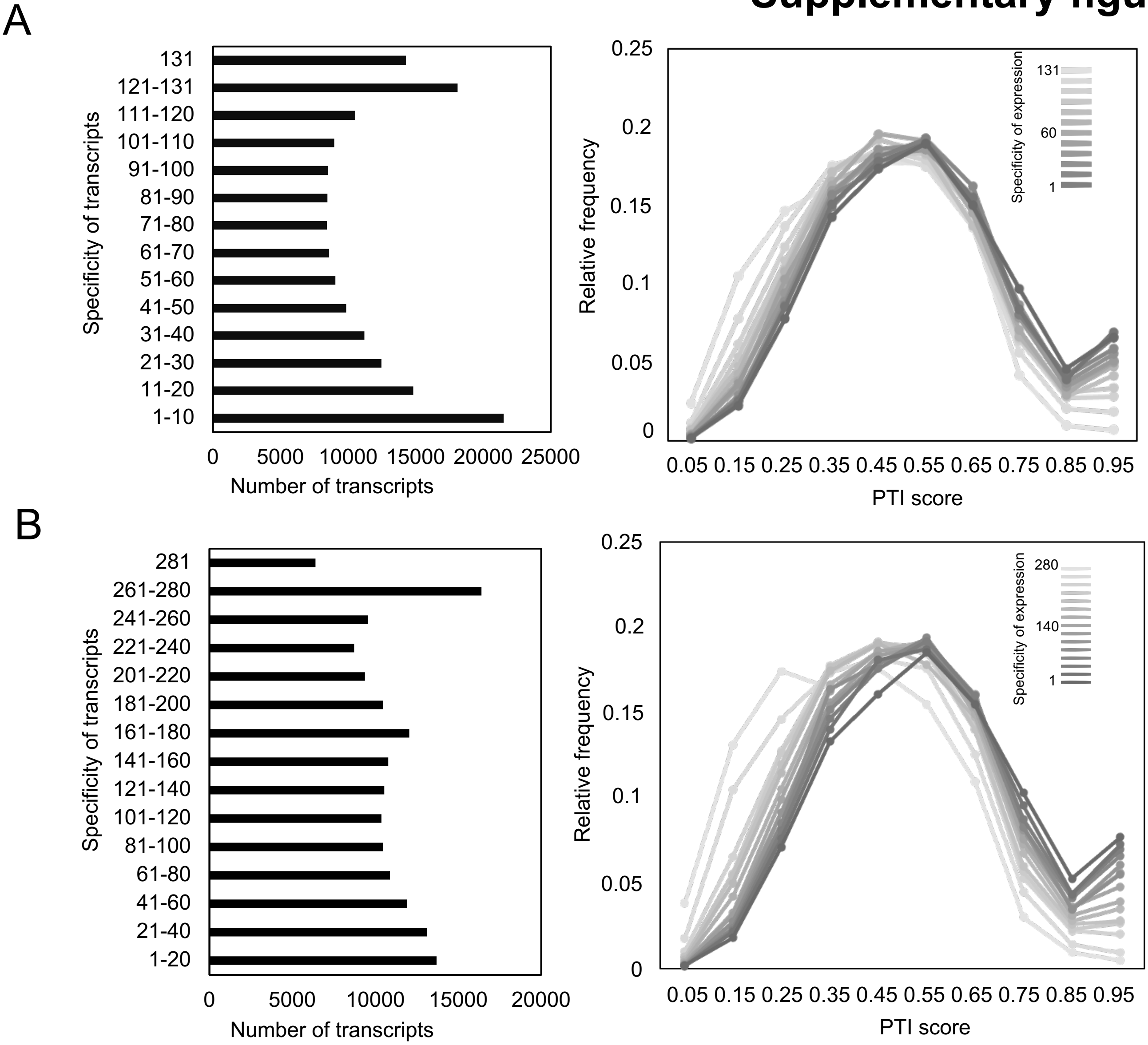
