## Supplementary Notes for "Protein-coding potential of RNAs measured by potentially translated island scores"

### Potentially translated islands

Sequences that begin at AUG and end at the 3′ terminus of RNA without UAA, UAG, or UGA are not considered PTIs. Hence, an RNA sequence lacking the AUG or the three base sequences that constitute stop codons (UAA, UAG, or UGA) does not contain PTIs. We did not use the reverse complement sequences of RNA sequences registered in databases to define PTIs because ribosomes translate mRNAs in the 5' to 3' direction.

### PTI score

In some transcripts, multiple PTIs have the longest length, causing the definition of pPTI and sPTI to become obscure owing to multiple PTIs with the same length. However, this is resolved by defining the PTI score only using the sum of all PTI lengths $l_{\mathrm{pPTI}} + \sum_{i=1}^{n} l_{\mathrm{sPTI}i}$ in the denominator, and the length of pPTI to calculate PTI score. Therefore, the PTI score is uniquely calculated even for those transcripts for which the pPTI cannot be clearly defined. If an RNA sequence does not have a PTI, both the numerator and denominator are set to 0. In such transcripts, there is no protein-coding potential, and the PTI score is not defined. Transcripts without PTIs were excluded from our analyses.
